# Chromatin Fiber Folding Represses Transcription and Loop Extrusion in Quiescent Cells

**DOI:** 10.1101/2020.11.24.396713

**Authors:** Sarah G. Swygert, Dejun Lin, Stephanie Portillo-Ledesma, Po-Yen Lin, Dakota R. Hunt, Cheng-Fu Kao, Tamar Schlick, William S. Noble, Toshio Tsukiyama

**Affiliations:** Basic Sciences Division, Fred Hutchinson Cancer Research Center, 1100 Fairview Ave. N, Seattle, WA 98109, USA; Department of Genome Sciences, University of Washington, 3720 15^th^ Ave. NE, Seattle, WA 98195, USA; Department of Chemistry, New York University, 100 Washington Square E, New York, NY 10003, USA; Institute of Cellular and Organismic Biology, Academia Sinica, 128 Sec. 2 Academia Road, Nankang, Taipei 11529, Taiwan; Courant Institute of Mathematical Sciences, New York University, 251 Mercer St, New York, NY, 10012, USA; New York University-East China Normal University Center for Computational Chemistry at New York University Shanghai, 3663 North Zhongshan Road, Shanghai, 200062, China; Paul G. Allen School of Computer Science and Engineering, University of Washington, 185 E Stevens Way NE, Seattle, WA 98195, USA

## Abstract

Determining the conformation of chromatin in cells at the nucleosome level and its relationship to cellular processes has been a central challenge in biology. We show that in quiescent yeast, widespread transcriptional repression coincides with the local compaction of chromatin fibers into structures that are less condensed and more heteromorphic than canonical 30-nanometer forms. Acetylation or substitution of H4 tail residues decompacts fibers and leads to global transcriptional de-repression. Fiber decompaction also increases the rate of loop extrusion by condensin. These findings establish a role for H4 tail-dependent local chromatin fiber folding in regulating transcription and loop extrusion in cells. They also demonstrate the physiological relevance of canonical chromatin fiber folding mechanisms even in the absence of regular 30-nanometer structures.

---

A longstanding hypothesis in biology is that chromatin fiber folding mediated by contacts among neighboring nucleosomes on the same DNA strand regulates DNA-dependent processes in eukaryotes. Although research over the past several decades has revealed mechanisms by which biochemically reconstituted chromatin fibers fold *in vitro*, determining three-dimensional chromatin structure at the nucleosome level in cells and its relation to cellular processes has been challenging (*1*, *2*). We previously used a genomics technique called Micro-C XL to map chromosomal interactions genome-wide at single-nucleosome resolution in purified quiescent *Saccharomyces cerevisiae* (*3*–*5*). Like their mammalian counterparts, quiescent budding yeast undergo dramatic chromatin condensation and widespread transcriptional repression (*6*–*8*), making them an ideal model for determining the mechanisms and functions of chromatin fiber folding in transcriptional regulation. We found that in quiescent yeast, the Structural Maintenance of Chromosomes (**SMC**) complex condensin relocates from its positions at tRNA genes, centromeres, and the rDNA in cycling (**log**) cells to form chromatin loop domains, termed large chromosomally interacting domains (**L-CIDs**), whose boundaries are at the promoters of coding genes (*3*). Condensin depletion during quiescence entry de-represses about 20% of all genes, with genes within 1 kilobase (**kb**) from a condensin binding site disproportionately represented. Although this represents a major form of transcriptional repression during quiescence, it does not account for the mechanism of repression of the majority of genes, which are located further within L-CID domains. We hypothesized that the folding of chromatin fibers within loops mediated by inter-nucleosomal interactions contributes to large-scale transcriptional repression in quiescent cells.

To examine differences between inter-nucleosomal interactions in log and quiescent cells, we used our previously published Micro-C XL data to generate maps of contact probabilities of nucleosomes within 1 kb of each other on the DNA strand (**Fig. 1A**) (*3*). Contact probability analysis revealed that the pattern of inter-nucleosomal interactions shifts between log and quiescence. In log, next-door neighbor *n+1* interactions are strongly favored, with interactions strongly decreasing with increasing distance. The high proportion of *n+1* interactions reflects an extended chromatin fiber where nucleosomes are most likely to encounter the nucleosomes closest to them on the DNA strand through random dynamics (*9*). In contrast, in quiescent cells, *n+1* and *n+2* contact counts are more similar, and longer-range interactions are more frequent. The propensity of nucleosomes to interact at greater distance in quiescent cells is consistent with a locally folded chromatin fiber. Biochemical experiments using reconstituted components and modeling studies have shown that canonically folded chromatin fibers with short linker DNA lengths form zig-zag 30 nanometer (**nm**) fiber structures with dominating *n+2* interactions (*10*–*14*). It was possible that lower than expected *n+2* interactions in the quiescent cell data might result from the Micro-C XL protocol, in which cells are crosslinked with both a short (formaldehyde) and a long crosslinker (disuccinimidyl glutarate [**DSG**]) (*5*). To test this, we repeated Micro-C XL of log and quiescent cells in the absence of DSG. As previously, and for all Micro-C XL experiments throughout, we used the HiCRep method to verify agreement between two biological replicates prior to merging replicate data to achieve maximum read depth, then used HiCRep to quantify the differences between conditions (**Fig. S1A,B**) (*15*, *16*). Omitting DSG only subtly affected Micro-C XL results in quiescent cells, with 200 base pair resolution (**bp**) (**Fig. 1B**), genome-wide (**Fig. S2A**), and 1 kb resolution (**Fig. S2B**) Micro-C XL heatmaps all appearing very similar. Consistently, the HiCRep stratum-adjusted correlation coefficient (**SCC**) calculated between quiescent cell Micro-C samples with and without DSG was 0.95 (**Fig. S1B**), the same as between biological replicates performed the same way (**Fig. S1A**). In contrast, omitting DSG from Micro-C XL of log cells diminished local interactions (**Fig. 1C**), though genome-wide and 1 kb resolution data appear more similar to each other (**Fig. S2C,D**). This suggests that use of long crosslinkers disproportionately captures longer-range and/or more transient local inter-nucleosomal interactions in log cells, consistent with a less folded and more dynamic chromatin fiber in log versus quiescent cells. Additionally, genome-wide metaplots generated +/− 10 kb from condensin-bound L-CID boundaries show that local inter-nucleosomal contacts in log cells occur at much lower distance and frequency than in quiescent cells (**Fig. 1D**, **S2E**). However, although local inter-nucleosomal interactions decreased overall when DSG was omitted, the pattern of nucleosome interactions in log and quiescent cell data remained similar to when DSG was included (**Fig. S2F**), confirming the original result. These data show that quiescent cell fibers are more compact than log fibers, but the compaction occurs via a different pattern of nucleosome interactions than predicted from biochemical experiments.

**Fig. 1.**
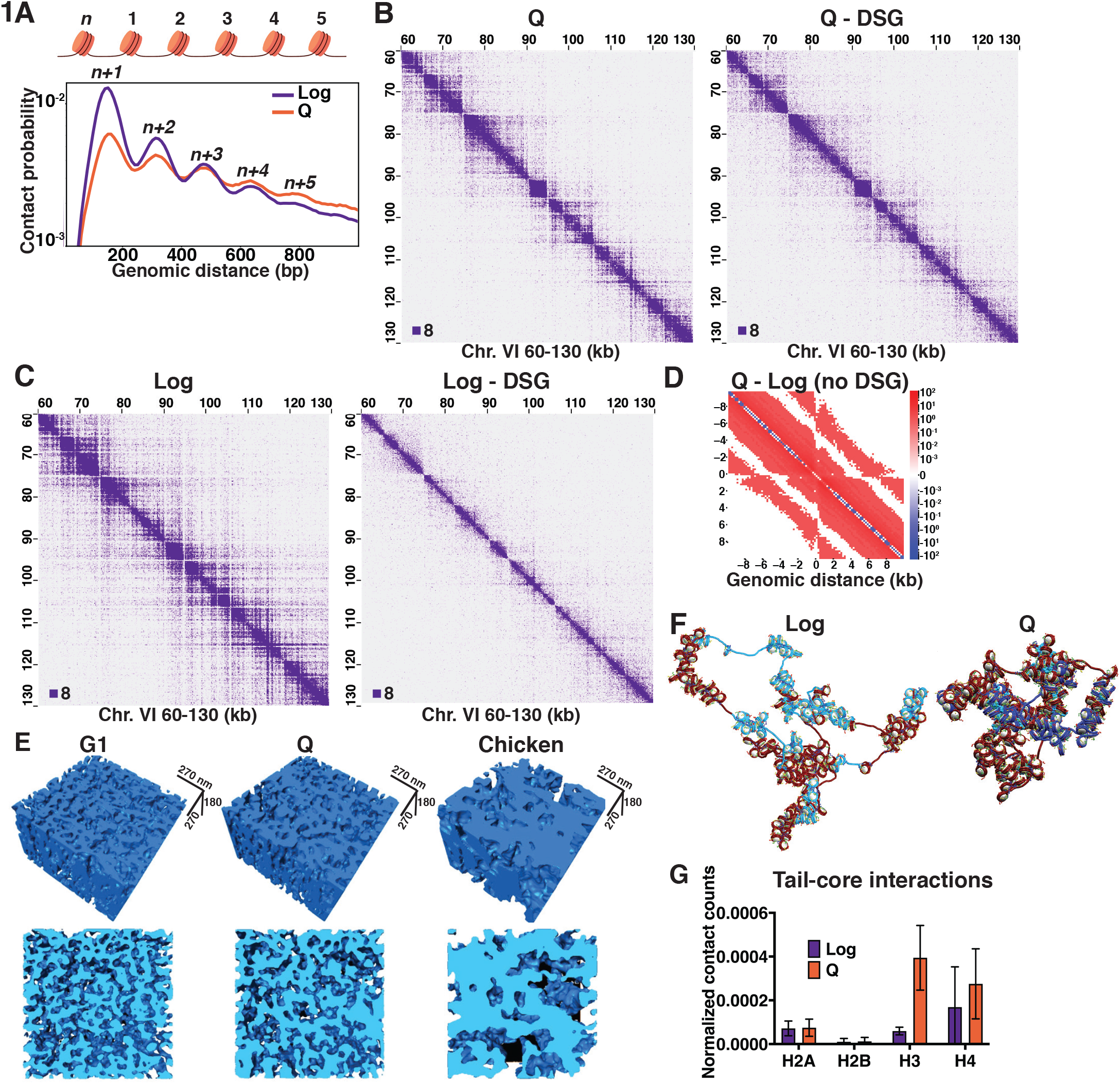
Chromatin fibers are folded in quiescent cells. (**A**) Contact probability map of nucleosome (*n*) interactions from Log and quiescent (Q) Micro-C XL data (*3*). Contacts between ligated di-nucleosomes in the “same” orientation (including “in-out” and “out-in” pairs) are shown (*4*). (**B**) Quiescent and (**C**) Log Micro-C XL with or without DSG at 200 bp resolution. For this and all subsequent Micro-C analyses, all in-facing read pairs were omitted to avoid contamination from undigested di-nucleosomes. (**D**) Subtraction Micro-C XL metaplots of median interactions +/−10 kb around sites of condensin-bound L-CID boundaries at 200 bp resolution. The scale shows difference between median counts. (**E**) Three-dimensional tomographic reconstructions of yeast and magnesium-treated chicken erythrocyte nuclei. (**F**) Representative equilibrated configurations of Log and Q chromatin fiber models of Chromosome I, 130-170 kb. Genes are shown in blue and intergenic regions are shown in red. Linker histone is shown in turquoise, and histone tails are shown in blue (H3), green (H4), yellow (H2A), and red (H2B). (**G**) Average frequency of histone tail/non-parental histone core contacts across 30 trajectories. Error bars show standard deviation.

To observe the conformation of folded quiescent chromatin fibers directly, we took a high-angle annular dark-field scanning transmission electron microscopy (**HAADF-STEM**) tomography approach to image 200 nm cross sections of uranyl acetate and lead citrate stained G1-arrested and quiescent yeast cells. As a positive control, we also imaged magnesium-treated chicken erythrocyte nuclei to test our ability to observe 30 nm fibers (*2*, *17*). STEM tomograms were reconstructed and analyzed to calculate fiber diameters (**Fig. 1E**, **S3A**). Our results are consistent with chromatin fiber compaction in quiescent versus log cells, with quiescent cell fibers appearing more compact qualitatively and demonstrating an upward shift in diameter as compared to log (**Fig. 1E**, **S3A,B**). However, quiescent cell fibers were not as compact as 30 nm fibers in chicken erythrocyte nuclei (**Fig. 1E**, **S3A,B**). These data are consistent with the Micro-C XL results and demonstrate that chromatin fibers adopt an extended, folded structure in quiescent cells that is distinct from and more irregular than canonical 30 nm fibers.

To examine quiescent chromatin fibers in more detail, we next used a nucleosome-resolution mesoscale modeling approach to investigate the structure of a 40 kb region of the right arm of Chromosome I during log and quiescence (*18*). The mesoscale model incorporates experimentally determined nucleosome positions, H4 tail acetylation, and putative linker histone occupancy (from MNase-seq, and histone H3, histone H4 tail penta-acetylation, and Hho1 ChIP-seq data) as developed recently to model genes (**Fig. S4A**) (*19*). Simulations of 30 independent trajectories converged after 80 million Monte Carlo steps (**Fig S4B,C**). Modeling revealed the same 40 kb region to be more compact in quiescent compared to log cells, with the radius of gyration decreasing and nucleosome packing increasing proportionally (**Fig. 1F**, **S4D-E**). The modeled contact probability also predicted increased *n*+1 contacts in log cells and a shift toward longer-range inter-nucleosomal interactions in quiescent cells, though more modest than in experimental data (**Fig. S4F**). Measurements were also more variable for simulations of the fiber in log, consistent with a more dynamic chromatin fiber in log versus quiescent cells (**Fig. S4E**). To investigate the mechanisms behind quiescent chromatin folding, we measured the frequency of contacts across configurations between histone tails and non-parental nucleosome linker DNA (**Fig. S4G**), histone tails and other histone tails (**Fig. S4H**), and histone tails and non-parental nucleosome cores (**Fig. 1G**). Although tail-nucleosome interactions were rare compared to tail-DNA and tail-tail interactions, H3 and H4 tails demonstrated a striking increase in non-parental nucleosome core interactions in quiescence compared to log.

Biochemical studies have shown that 30 nm fiber folding depends on a basic patch in the histone H4 tail, which at least in part mediates folding by making contacts with an acidic patch on the surface of a nearby nucleosome (*10*, *20*–*23*). This folding is disrupted by acetylation of the H4 basic patch at lysine 16 (*24*). One of the primary differences between log and quiescent chromatin is the massive deacetylation of residues in the histone H3 and H4 tails during quiescence entry due to global targeting of the histone deacetylase complex Rpd3 (*25*). Because of this and our modeling data showing an increase in histone tail-nucleosome core interactions, we wondered if chromatin folding in quiescent cells may function by a similar mechanism as 30 nm fiber folding. To test this, we treated cells during a late stage of quiescence entry with the histone deacetylase inhibitor Trichostatin A (**TSA**), which has previously been shown to disrupt inter-nucleosomal interactions in mammalian cells (*26*, *27*). TSA treatment of quiescent cells led to an increase in H4 tail acetylation approaching log cell levels as measured by ChIP-seq (**Fig. 2A**), and higher than log in bulk as measured by Western blot (**Fig. S5A**). To determine the effect of histone acetylation on global chromatin condensation in TSA-treated cells, we measured chromatin volume by staining DNA with 4′,6-diamidino-2-phenylindole (**DAPI**) followed by confocal imaging. TSA treatment significantly increased the chromatin volume of quiescent cells (**Fig. 2B**, **S5B**). To examine if this change resulted from alterations in local chromatin folding, we next performed Micro-C XL on TSA-treated quiescent cells. Micro-C XL data at 200 bp resolution show that TSA treatment does reduce local chromatin interactions (**Fig. 2C**; see **Fig. S5C-D** for examples at lower resolution). Consistently, the pattern of intra-nucleosomal interactions shifted so that *n+1* and *n+2* interactions increased and longer-range interactions decreased (**Fig. S5E**).

**Fig. 2.**
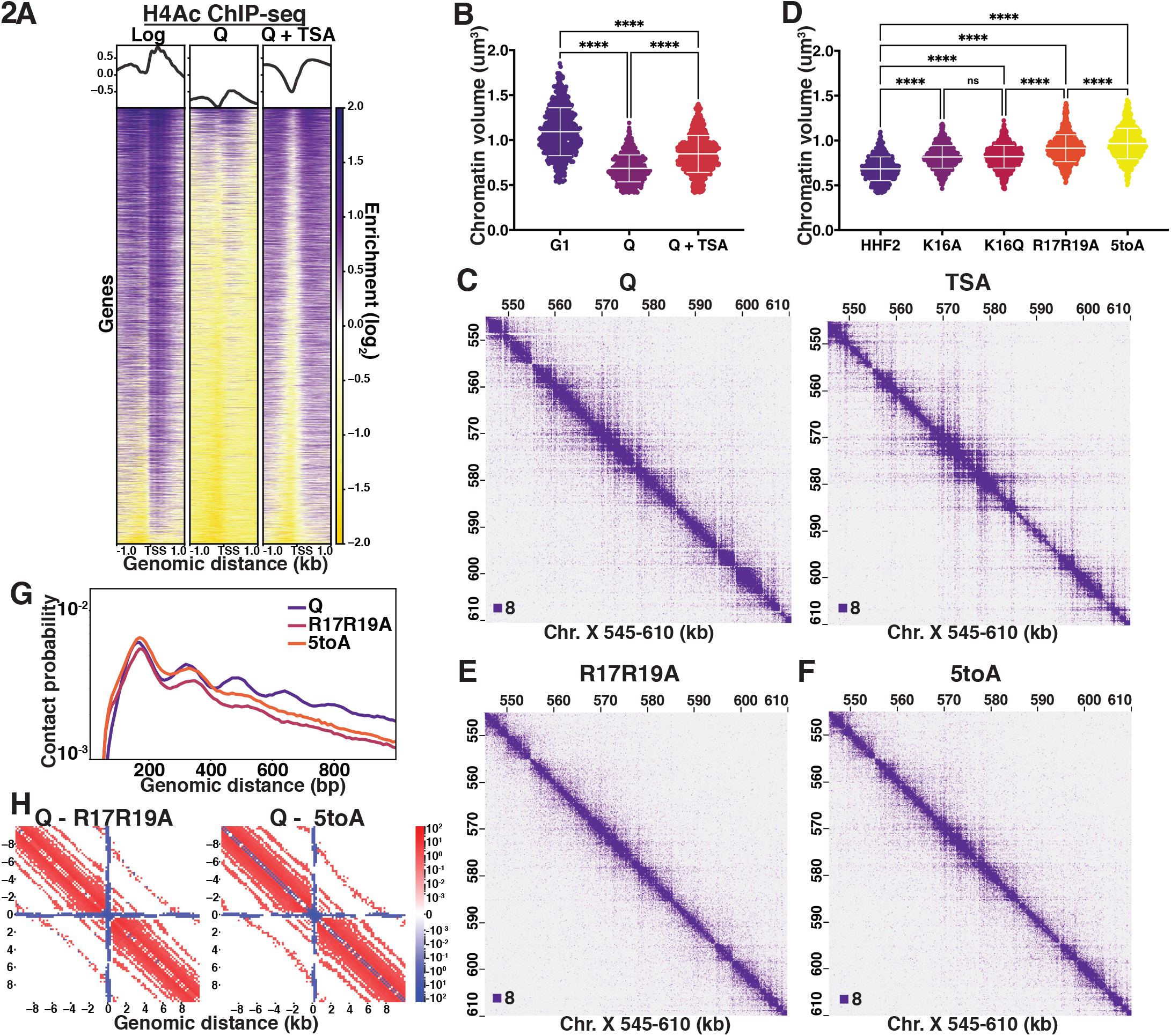
The H4 tail basic patch regulates chromatin folding in quiescent cells. (**A**) H4 tail penta-acetylation ChIP-seq heatmaps +/−1 kb of all transcription start sites (TSS). Rows are linked across all heatmaps. (**B**) Chromatin volume measurements following DAPI staining of at least 200 cells each of two biological replicates. Bars represent mean and standard deviation. Significance was determined using two-tailed paired t-tests. (**C**) Micro-C XL data of quiescent cells without (left) and with (right) TSA treatment at 200 bp resolution. (**D**) Volume measurements as in (B) following DAPI staining of quiescent H4 mutant cells. HHF2 is the single-copy HHT2-HHF2 WT control strain. (**E**) and (**F**) Micro-C XL data of H4 mutant quiescent cells at 200 bp resolution. (**G**) Contact probability map generated from Micro-C XL data. Q indicates quiescent cells from true WT strains. (**H**) Subtraction Micro-C XL metaplots of median interactions around sites of condensin-bound L-CID boundaries in quiescent cells at 200 bp resolution. The scale shows difference between median counts.

These data support histone tail deacetylation as being crucial to quiescent chromatin fiber folding at the local scale. However, while the H4 basic patch consists of five residues, only two, lysines 16 (**K16**) and 20 (**K20**), are capable of being acetylated. To test the involvement of all five H4 tail basic patch residues, we created yeast strains in which the endogenous H3 and H4 loci were deleted and complemented by a mutant or wild type copy of H3 and H4 genes at an ectopic locus. The wild type control strains (**HHF2**) grow and enter quiescence very similarly to true wild type (**WT**) strains with two copies of H3 and H4 genes. DAPI staining and volume measurements of quiescent cells show that alanine and glutamine substitutions of K16 (**K16A** and **K16Q**) significantly increase quiescent cell chromatin volume to a similar extent (**Fig. 2D**, **S6A**). Substitution of both arginine residues with alanine (**R17R19A**) decompacts chromatin even further, and full abrogation of all five basic patch residues (^16^KRHRK^20^) with alanine (**5toA**) decompacts chromatin almost to G1 levels. Consistent with the basic patch playing an important role in quiescence, H4 strains bearing basic patch substitutions do not display altered growth in log, but have up to an 80% reduction in quiescence entry (**Fig. S6B**), although cells that did enter maintained similar longevity to WT (**Fig. S6C**). Substitutions to residues in the acidic patch similarly increased chromatin volume (**Fig. S6D**), supporting the model that H4 tail/acidic patch contacts may be involved in chromatin folding in quiescent cells. However, these H2A mutant strains exhibited stronger growth defects and were more difficult to work with than H4 mutants, and we did not pursue further experiments with them.

We next completed Micro-C XL experiments using quiescent cells of the mutants displaying the most chromatin decompaction, R17R19A and 5toA, as well as the control HHF2, which was largely indistinguishable from WT (**Fig. S1B**). In contrast, both basic patch mutants displayed strong chromatin fiber decompaction at the local level in quiescent cells (**Fig. 2E,F**). Other than telomere clustering defects evident in genome-wide plots as previously shown (*28*), longer-range interactions displayed less prominent differences from WT (**Fig. S6E-H**). Consistent with massive chromatin unfolding, the pattern of local inter-nucleosomal interactions dramatically shifted, with interactions beyond *n+2* strongly decreasing in both mutants, and *n+1* interactions increasing relative to *n+2* (**Fig. 2G**). Metaplots centered around L-CID boundaries similarly showed a large reduction in local chromatin contacts as compared to WT (**Fig. 2H**, **S6I**). Collectively, these experiments support a model in which chromatin fiber folding in quiescent cells is mediated by inter-nucleosomal interactions driven by the H4 basic patch.

We next sought to determine the role of H4 tail-mediated chromatin fiber folding in transcriptional repression during quiescence. To this end, we completed ChIP-seq of the RNA Polymerase II (**Pol II**) subunit Rpb3 in log and quiescent mutant cells. We have previously shown that Rpb3 ChIP-seq correlates well with ChIP-seq of phosphorylated serine 2 of the Pol II carboxy-terminal domain and is thus a good way of measuring active transcription (*3*, *29*), while avoiding complications in measurements introduced by the storage of mRNAs in stress granules (*3*, *30*) and processing bodies (*31*) during quiescence. Consistent with previous results (*32*), genome-browser tracks (**Fig. S7A**), and Pearson correlation scores (**Fig. S7B**) displayed minimal differences in Pol II occupancy between basic patch mutants and HHF2 in log. In contrast, Pol II occupancy increased dramatically in quiescent basic patch mutant cells, with occupancy increasing proportionally to the number of substitutions introduced to the basic patch (**Fig. 3A-C**, **S7A, S7C**). ChIP-seq peak calling implemented using MACS2 similarly showed a dramatic increase in Pol II peaks between HHF2 and 5toA quiescent cells, with the number of peaks increasing in the 5toA mutant corresponding to approximately 40% of all genes (**Fig. 3D**).

**Fig. 3.**
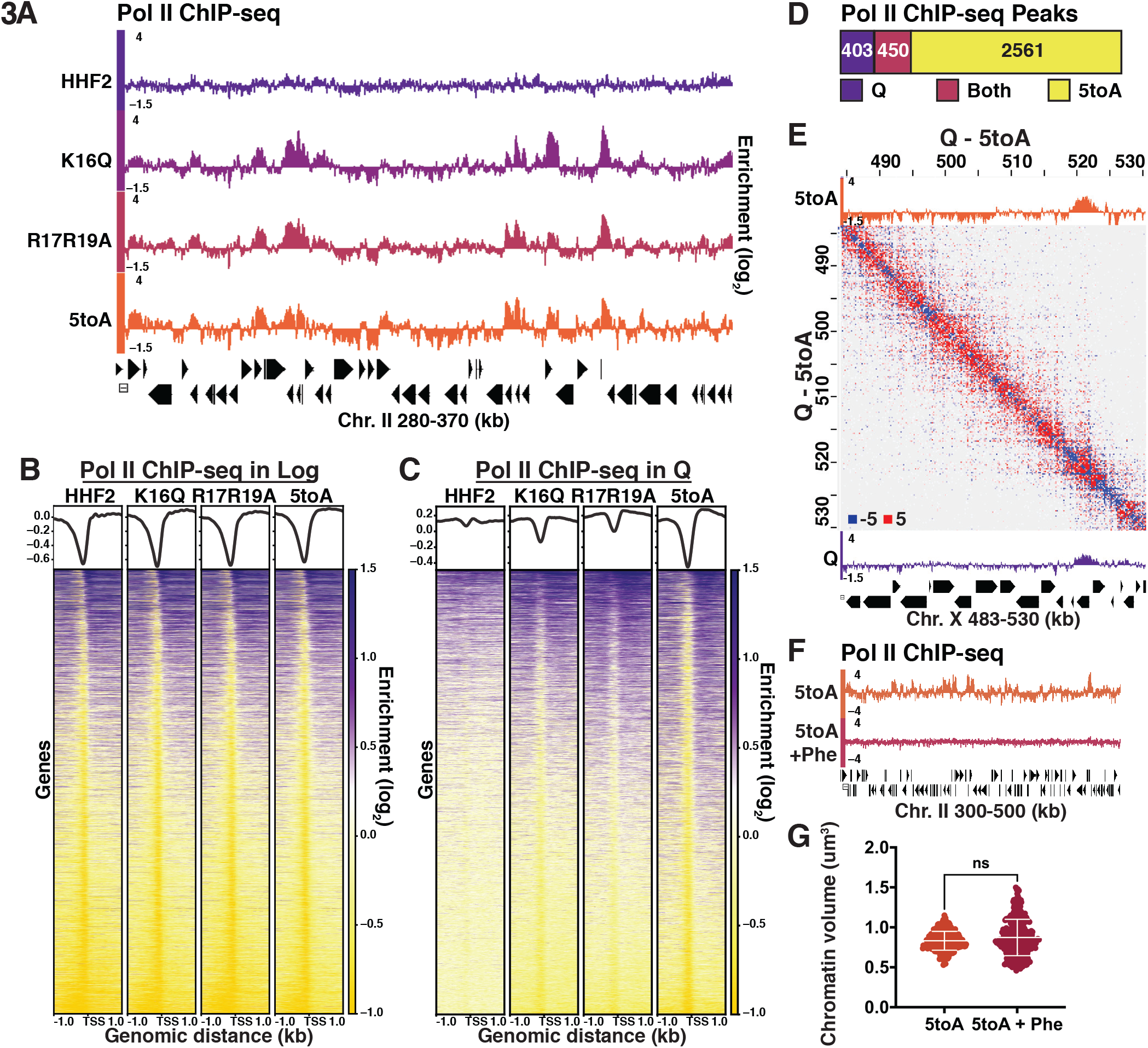
H4 tail-mediated chromatin folding represses transcription during quiescence. (**A**) Genome browser view of Pol II subunit Rpb3 ChIP-seq data in quiescent mutant strains across a portion of Chromosome II. (**B**) Heatmaps of Rpb3 across all TSSs in Log and (**C**) Q cells. Rows are the same across all heatmaps in a panel. (**D**) MACS differential peak calls for WT and 5toA Rpb3 ChIP-seq in Q. (**E**) Pol II ChIP-seq data in 5toA (top) and true WT Q (bottom) cells overlaid on a heatmap showing 5toA Micro-C XL data subtracted from WT Q data. Positive (red) indicates contacts that are higher in WT cells. (**F**) Genome browser view of Rpb3 ChIP-seq data in 5toA quiescent cells with or without 1,10-phenanthroline treatment (5toA+Phe). (**G**) Chromatin volume measurements following DAPI staining of at least 200 cells each of two biological replicates. Bars represent mean and standard deviation. Significance was determined using two-tailed paired t-tests.

Although these results strongly support the model that local chromatin fiber folding mediated by the H4 tail represses global transcription in quiescent cells, another possible interpretation of these data is that transcriptional activation leads to chromatin fiber decompaction. However, activation appears to be epistatic to chromatin decompaction, as fiber unfolding occurs in the basic patch mutants genome-wide, regardless of gene expression level (**Fig. 3E**, **S7D-F**). This suggests that chromatin folding represses transcription, but unfolding is not sufficient for activation. To test this idea more directly, we treated 5toA cells with the transcription inhibitor 1,10-phenanthroline during quiescence entry. Phenanthroline treatment led to complete loss of Pol II occupancy by ChIP-seq (**Fig. 3F**). However, DAPI staining of phenanthroline-treated 5toA quiescent cells showed a slight non-significant increase in chromatin volume (**Fig. 3G**), supporting that chromatin decondensation in 5toA cells is a cause and not a result of global transcriptional activation.

Unexpectedly, upon examination of the Micro-C XL data from basic patch mutant quiescent cells, we also observed the striking appearance of stripes that overlapped with condensin subunit Brn1 ChIP-seq peaks in WT quiescent cells (**Fig. 4A**, **2H**, **S8A-C**). Stripes are believed to occur in Hi-C data as a result of loop extrusion, in which SMC complexes bind the genome at two points and increase the size of the resulting loop by moving chromatin in one or both directions through the complex (*33*–*35*). In the case of one-sided extrusion, chromatin at the fixed boundary progressively contacts chromatin that is being extruded, leading to the formation of a stripe when contacts are measured at the population level (**Fig. 4B**). The SMC complex would also be expected to bind extruded chromatin progressively. Consequently, this result suggests that substitutions in the H4 tail affect the way in which condensin extrudes chromatin loops. To determine if this change is also reflected in condensin localization, we performed ChIP-seq of Brn1-FLAG in the mutants. Although stripes are not observable at all Brn1 sites, metaplots generated from WT, R17R19A, and 5toA Micro-C data showing *trans* interactions between Brn1 peaks show clear stripe patterns at the median in the mutants (**Fig. 4C**). Although bulk Brn1 protein levels did not appreciably change between the mutants and HHF2 (**Fig. S8D**), Brn1 localization was altered in the mutants (**Fig. 4A,D** and **S8E**). While Brn1 shows strong localization at nucleosome-depleted regions in WT and HHF2 quiescent cells, basic patch mutants show a reduction in the magnitude of Brn1 peaks and a corresponding flattening out across gene bodies surrounding promoters (**Fig. 4D**) and L-CID boundaries (**Fig. 4E**). MACS peak calling further showed a decrease in the number of Brn1 peaks between HHF2 and 5toA (**Fig. 4F**), which we interpret to mean that Brn1 spreading across the genome reduces many discrete peaks to below detectable levels. Overlaying Brn1 ChIP-seq tracks on the Micro-C XL data shows that stripes overlap well with Brn1 sites in the mutants, and that the broadening of Brn1 peaks is easily observable at the level of individual loci (**Fig. 4A**, **S8A-C**). Importantly, this flattening and spreading of Brn1 signals is consistent with the stripes observed in the Micro-C XL data, with the strongest stripe interactions emanating immediately away from L-CID boundaries and then obvious loops forming between Brn1 peaks at L-CID boundaries (**Fig. 4A,C**). This shows that stripes at least in part result from progressive Brn1 binding of regions of chromatin within L-CIDs, as would be expected from active loop extrusion.

**Fig. 4.**
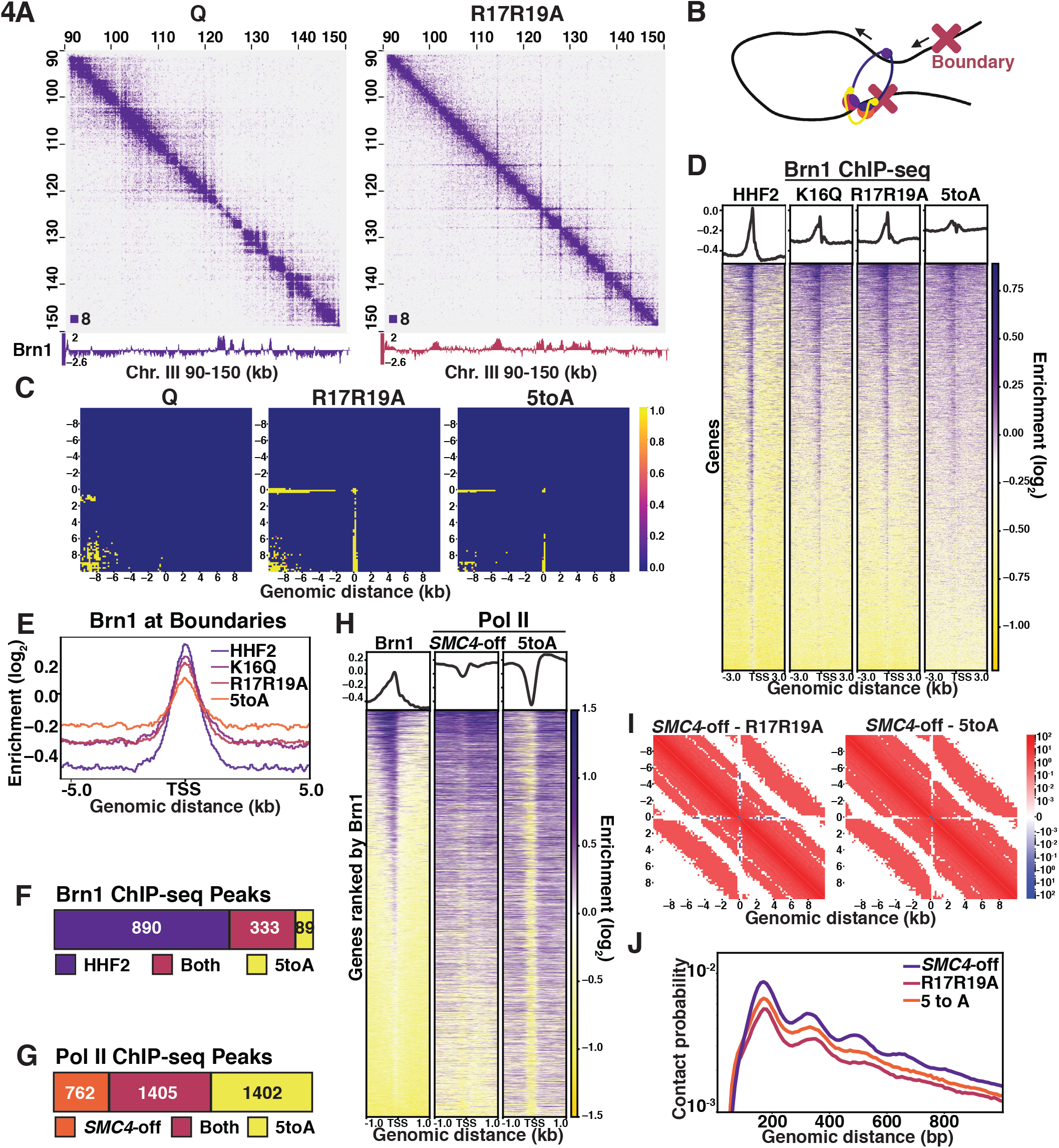
H4-tail mediated chromatin folding inhibits condensin loop extrusion. (**A**) Condensin subunit Brn1 ChIP-seq data overlayed beneath Micro-C XL data at 200bp resolution. (**B**) Schematic showing one-sided loop extrusion by condensin between two boundaries. (**C**) Metaplots of Micro-C XL data showing *trans* nucleosome contacts +/−10kb between Brn1 ChIP-seq peaks in the indicated condition. (**D**) Heatmaps of Brn1 ChIP-seq +/−3kb of all TSSs. Rows are the same across all heatmaps. (**E**) Metaplot of Brn1 ChIP-seq data +/−5kb of L-CID boundaries. (**F**) MACS differential peak calls for HHF2 and 5toA Brn1 ChIP-seq in Q. (**G**) MACS differential peak calls for *SMC4*-off and 5toA Brn1 ChIP-seq in Q. *SMC4*-off cells contain a doxycycline-inducible Tet repressor system to shut off expression of the condensin subunit SMC4. *SMC4*-off data were previously published in (*3*). (**H**) Heatmaps of Brn1 and Pol II (subunit Rbp3) ChIP-seq +/−1kb of all TSSs. All heatmaps are ordered by descending Brn1 occupancy in WT Q (leftmost heatmap). (**I**) Micro-C XL subtraction metaplots of median interactions around sites of condensin-bound L-CID boundaries in quiescent cells at 200 bp resolution. The scale shows difference between median counts. Plots show mutant Micro-C XL data subtracted from previously published *SMC4*-off Micro-C XL data (*3*). (**J**) Contact probability map of Micro-C XL data.

Modeling studies have shown that the emergence of stripes in Hi-C data is likely to result from an increase in the rate at which SMC complexes extrude chromatin loops: as loop extrusion occurs more rapidly, interacting regions of chromatin have less time to diffuse apart and consequently retain “memory” of being extruded (*36*). Additionally, constant rapid extrusion is likely to be captured at all points of passage at the cell population level. Although transcription has been proposed to affect loop extrusion through multiple avenues — by slowing the process of extrusion by creating difficult-to-traverse transcription bubbles, by “pushing” SMC complexes across chromatin, and by promoting SMC complex loading — we do not believe the change in condensin extrusion that we see in the H4 mutants is due to the increase in transcription in the mutants (*37*–*40*). This is because 1) stripes occur uniformly between L-CID boundaries, regardless of Pol II occupancy (as seen in **Fig. 4A**, **S8A-C**); 2) stripes appear as strong in the R17R19A mutant as in the 5toA mutant, despite less Pol II occupancy overall; and 3) regions displaying stripes do not exhibit large changes in Pol II occupancy between WT and the mutants, because Brn1 sites tend to overlap with Pol II-occupied genes even in WT (**Fig. S8F**) (*3*). Instead, we propose a model in which H4 tail-mediated chromatin fiber folding sterically impedes the ability of condensin to extrude chromatin, leading to slower loop extrusion than in mutants with decompacted fibers. This slower extrusion may help to stabilize loops between L-CID boundaries in WT quiescent cells. We believe this is a complementary rather than primary mechanism of transcriptional repression by the H4 tail. While many Pol II peaks appear in both condensin subunit SMC4-depleted *SMC4*-off cells and the 5toA mutant, there are large numbers of non-overlapping Pol II peaks, with nearly double the number of distinct peaks in the 5toA mutant (**Fig. 4G,H**). As previously shown, genes with the highest Pol II occupancy in *SMC4*-off cells tend to be close to Brn1 sites, while Pol II occupancy in 5toA cells does not necessarily overlap with Brn1 (**Fig. 4H**). Additionally, although condensin depletion disrupts repression by decreasing insulation at L-CID boundaries and decreasing local inter-nucleosomal interactions to some extent compared to WT, the H4 mutants display significantly more decompaction than *SMC4*-off cells, while retaining some insulation at boundaries (**Fig. 4I,J**).

Although 30 nm chromatin fibers form robustly in biochemical experiments, they have been largely absent from studies examining chromatin fibers in cells, suggesting they tend to occur in a handful of specialized cell types (*41*). However, the expectation that chromatin fiber conformation regulates DNA-dependent processes persists both in textbooks and in the literature. While differences in fiber diameter have been observed between euchromatin and heterochromatin (*42*), and chromatin nanodomain cluster size was found to increase during B-cell activation (*43*), genome-wide changes in chromatin fiber conformation at the nucleosome level have not previously been observed in cells, even between interphase and mitotic chromosomes (*2*). Our results demonstrate that chromatin fibers adopt a dramatically different conformation during quiescence. These structures are less compact and more heteromorphic than canonical 30 nm fibers; however, their dependence on the H4 basic patch suggests they form through a similar mechanism. This finding demonstrates that mechanisms uncovered through studies of biochemically reconstituted chromatin fibers have physiological relevance despite the irregularity of *in vivo* fibers. Additionally, although differences in the length of short clutches of interacting nucleosomes have been observed in active versus repressed genes (*4*, *26*, *44*), changes in chromatin fiber structure have not previously been shown to regulate transcription. Our data show that chromatin fiber folding acts strongly to repress transcription in cells. They also suggest that the conformation of the underlying chromatin fiber plays a role in the progression of loop extrusion by SMC complexes. As nucleosome-resolution chromatin tools become more widely available, we expect that local chromatin compaction will be further demonstrated to regulate processes in a variety of contexts and cell types.

## Acknowledgments

The authors would like to thank the past and current members of the Tsukiyama lab, especially Tianhong Fu, Laura Hsieh, and Jeffrey McKnight. We would also like to thank the Fred Hutch Genomics shared resource. Portions of some figures were created with BioRender.com. We apologize to colleagues whose work we were unable to cite due to space constraints.

## Author contributions

Conceptualization, all authors; Methodology, all authors; Software, D.L. (lead), S.G.S., S.P.-L.; Validation, S.G.S., D.L., S.P.-L, P.-Y.L., D.R.H.; Formal Analysis, D.L. (lead), S.G.S., S.P.-L, P.-Y.L., D.R.H.; Investigation, S.G.S. (lead), S.P.-L, P.-Y.L., D.R.H.; Resources, S.G.S., D.R.H., C.-F.K., T.T.; Data Curation, S.G.S., D.L.; Writing – Original Draft, S.G.S. (lead), T.T. (lead), D.L., S.P.-L, P.-Y.L., D.R.H.; Writing – Review & Editing, all authors; Visualization, S.G.S., D.L., S.P.-L, P.-Y.L., D.R.H.; Supervision, T.S., W.S.N., T.T.; Project Administration, S.G.S., T.T.; Funding Acquisition, S.G.S., P.-Y.L., C.-F.K., T.S., W.S.N., T.T.

## Data and materials availability

Genomics data will be made publicly accessible on NCBI following acceptance of publication. Plasmids and yeast strains are available on request.

## Supplementary Materials

## Materials and Methods

### Yeast growth and quiescent cell purification

Quiescent cells were grown as previously described (*6*, *45*), by using single colonies to inoculate cultures in rich media in flasks with at least a 1:5 ratio of culture to flask volume. Cells were grown for seven days, pelleted, and resuspended in 2.5 mL of water prior to loading on density gradients. Gradients were prepared using 25 mL of a mixture containing 90% Percoll (GE, catalog #17-0891-01) and 150 mM NaCl in 50 mL high-speed round bottom centrifuge tubes. Gradients were centrifuged at 10,000 g for 15 min prior to loading samples, then centrifuged at 300 g for 1 hour. Quiescent cells were removed from the bottom of gradients by pipetting, washed in water, and quantified using spectrophotometry. For TSA-treatment, TSA (Fisher, catalog # 58880-19-6) was resuspended to 50 mM in DMSO, and added to cultures to 50 μM 24 hours prior to purifying quiescent cells. During purification, 50 μM TSA was also added to Percoll gradients and water used to resuspend cells. For 1,10-phenanthroline treatment, phenanthroline was dissolved in methanol to 100 mg/mL, then added to cultures for a final concentration of 150 ug/mL 24 hours prior to purifying quiescent cells. For G1-arrested cells, cultures were inoculated to an optical density at A_660nm_ of 0.06, then α-factor was added to 10 μg/mL once the optical density reached 0.15. Cultures were monitored for G1 arrest under the microscope until at least 95% appeared to be in G1, approximately 90 minutes later.

### Yeast strains

| Strain | Source | Identifier |
| --- | --- | --- |
| WT: MATa RAD5+ prototroph | (25) | yTT5781 |
| WT: MATa RAD5+ prototroph | (25) | yTT5783 |
| Hho1-FLAG: MATa RAD5+ prototroph HHO1-2L-3FLAG::KanMX | This Paper | yTT6336 |
| Hho1-FLAG: MATa RAD5+ prototroph HHO1-2L-3FLAG::KanMX | This Paper | yTT6337 |
| HHF2: MATa RAD5+ ura3-1 hht1-hhf1::Nat hht2-hhf2::Hyg trp1-1::pRS404-HHT2-HHF2 | This Paper | yTT7177 |
| HHF2: MATa RAD5+ ura3-1 hht1-hhf1::Nat hht2-hhf2::Hyg trp1-1::pRS404-HHT2-HHF2 | This Paper | yTT7206 |
| hhf2-5toA: MATa RAD5+ ura3-1 hht1-hhf1::Nat hht2-hhf2::Hyg trp1-1::pRS404-HHT2-hhf2-K16A,R17A,H18A,R19A,K20A | This Paper | yTT7175 |
| hhf2-5toA: MATa RAD5+ ura3-1 hht1-hhf1::Nat hht2-hhf2::Hyg trp1-1::pRS404-HHT2-hhf2-K16A,R17A,H18A,R19A,K20A | This Paper | yTT7208 |
| hhf2-R17A,R19A: MATa RAD5+ ura3-1 hht1-hhf1::Nat hht2-hhf2::Hyg trp1-1::pRS404-HHT2-hhf2-R17A,R19A | This Paper | yTT7200 |
| hhf2-R17A,R19A: MATa RAD5+ ura3-1 hht1-hhf1::Nat hht2-hhf2::Hyg trp1-1::pRS404-HHT2-hhf2-R17A,R19A | This Paper | yTT7207 |
| hhf2-K16A: MATa RAD5+ ura3-1 hht1-hhf1::Nat hht2-hhf2::Hyg trp1-1::pRS404-HHT2-hhf2-K16A | This Paper | yTT7202 |
| hhf2-K16A: MATa RAD5+ ura3-1 hht1-hhf1::Nat hht2-hhf2::Hyg trp1-1::pRS404-HHT2-hhf2-K16A | This Paper | yTT7209 |
| hhf2-K16Q: MATa RAD5+ ura3-1 hht1-hhf1::Nat hht2-hhf2::Hyg trp1-1::pRS404-HHT2-hhf2-K16Q | This Paper | yTT7205 |
| hhf2-K16Q: MATa RAD5+ ura3-1 hht1-hhf1::Nat hht2-hhf2::Hyg trp1-1::pRS404-HHT2-hhf2-K16Q | This Paper | yTT7210 |
| HHF2 Brn1-FLAG: MATa RAD5+ ura3-1 hht1-hhf1::Nat hht2-hhf2::Hyg trp1-1::pRS404-HHT2-HHF2 Brn1-2L-3FLAG::KanMX | This Paper | yTT7390 |
| **HHF2 Brn1-FLAG:** MATa RAD5+ ura3-1 hht1-hhf1::Nat hht2-hhf2::Hyg trp1-1::pRS404-HHT2-HHF2 Brn1-2L-3FLAG::KanMX | This Paper | yTT7391 |
| **hhf2-5toA Brn1-FLAG:** MATa RAD5+ ura3-1 hht1-hhf1::Nat hht2-hhf2::Hyg trp1-1::pRS404-HHT2-hhf2-K16A,R17A,H18A,R19A,K20A Brn1-2L-3FLAG::KanMX | This Paper | yTT7388 |
| **hhf2-5toA Brn1-FLAG:** MATa RAD5+ ura3-1 hht1-hhf1::Nat hht2-hhf2::Hyg trp1-1::pRS404-HHT2-hhf2-K16A,R17A,H18A,R19A,K20A Brn1-2L-3FLAG::KanMX | This Paper | yTT7389 |
| **hhf2-R17A,R19A Brn1-FLAG:** MATa RAD5+ ura3-1 hht1-hhf1::Nat hht2-hhf2::Hyg trp1-1::pRS404-HHT2-hhf2-R17A,R19A Brn1-2L-3FLAG::KanMX | This Paper | yTT7392 |
| **hhf2-R17A,R19A Brn1-FLAG:** MATa RAD5+ ura3-1 hht1-hhf1::Nat hht2-hhf2::Hyg trp1-1::pRS404-HHT2-hhf2-R17A,R19A Brn1-2L-3FLAG::KanMX | This Paper | yTT7393 |
| **hhf2-K16Q Brn1-FLAG:** MATa RAD5+ ura3-1 hht1-hhf1::Nat hht2-hhf2::Hyg trp1-1::pRS404-HHT2-hhf2-K16Q Brn1-2L-3FLAG::KanMX | This Paper | yTT7394 |
| **hhf2-K16Q Brn1-FLAG:** MATa RAD5+ ura3-1 hht1-hhf1::Nat hht2-hhf2::Hyg trp1-1::pRS404-HHT2-hhf2-K16Q Brn1-2L-3FLAG::KanMX | This Paper | yTT7395 |
| **HTA1:** MATa W303 Rad5+ ade2-1 can1-100 his3-11,15 leu2-3,112 ura3-1 hta1-htb1::Hyg hta2-htb2::Nat trp1-1::pRS404-HTA1-HTB1 | This Paper | yTT6767 |
| **HTA1:** MATa W303 Rad5+ ade2-1 can1-100 his3-11,15 leu2-3,112 ura3-1 hta1-htb1::Hyg hta2-htb2::Nat trp1-1::pRS404-HTA1-HTB1 | This Paper | yTT6773 |
| **hta1-E57A:** MATa W303 Rad5+ ade2-1 can1-100 his3-11,15 leu2-3,112 ura3-1 hta1-htb1::Hyg hta2-htb2::Nat trp1-1::pRS404-hta1-E57A-HTB1 | This Paper | yTT6768 |
| **hta1-E57A:** MATa W303 Rad5+ ade2-1 can1-100 his3-11,15 leu2-3,112 ura3-1 hta1-htb1::Hyg hta2-htb2::Nat trp1-1::pRS404-hta1-E57A-HTB1 | This Paper | yTT6774 |
| **hta1-E65A:** MATa W303 Rad5+ ade2-1 can1-100 his3-11,15 leu2-3,112 ura3-1 hta1-htb1::Hyg hta2-htb2::Nat trp1-1::pRS404-hta1-E65A-HTB1 | This Paper | yTT6769 |
| **hta1-E65A:** MATa W303 Rad5+ ade2-1 can1-100 his3-11,15 leu2-3,112 ura3-1 hta1-htb1::Hyg hta2-htb2::Nat trp1-1::pRS404-hta1-E65A-HTB1 | This Paper | yTT6776 |
| **hta1-E93A:** MATa W303 Rad5+ ade2-1 can1-100 his3-11,15 leu2-3,112 ura3-1 hta1-htb1::Hyg hta2-htb2::Nat trp1-1::pRS404-hta1-E93A-HTB1 | This Paper | yTT6772 |
| **hta1-E93A:** MATa W303 Rad5+ ade2-1 can1-100 his3-11,15 leu2-3,112 ura3-1 hta1-htb1::Hyg hta2-htb2::Nat trp1-1::pRS404-hta1-E93A-HTB1 | This Paper | yTT6779 |
| **hta1-E65A,D92A:** MATa W303 Rad5+ ade2-1 can1-100 his3-11,15 leu2-3,112 ura3-1 hta1-htb1::Hyg hta2-htb2::Nat trp1-1::pRS404-hta1-E65A,D92A-HTB1 | This Paper | yTT6618 |
| **hta1-E65A,D92A:** MATa W303 Rad5+ ade2-1 can1-100 his3-11,15 leu2-3,112 ura3-1 hta1-htb1::Hyg hta2-htb2::Nat trp1-1::pRS404-hta1-E65A,D92A-HTB1 | This Paper | yTT6765 |

### Micro-C XL

For quiescent cells, 2400 optical density units of purified cells were resuspended in 1.2 L water and crosslinked using 106.8 mL 37% formaldehyde for 10 min at 30°C with shaking. Formaldehyde was attenuated with 120 mL 2.5 M glycine, and cells were pelleted, washed, and resuspended in 120 mL Buffer Z (1 M sorbitol, 50 mM Tris pH 7.4, 10 mM β-mercaptoethanol). Cells were split into twelve 15 mL conical tubes and spheroplasted by addition of 1 mL 10 mg/mL 100T zymolyase at 30°C with rotation until at least 60% of cells appeared as spheroplasts under the microscope (approximately 45-120 minutes). Spheroplasts were centrifuged for 10 minutes at 3,000 rpm and 4°C, washed in cold PBS, and pelleted again. For DSG crosslinking, DSG (Fisher, #PI20593) was resuspendend to 300 mM in DMSO and diluted to 3 mM in room temperature PBS. Spheroplasts were resuspended in 5 mL DSG solution and crosslinked for 40 min at 30°C with rotation. Crosslinking was quenched by addition of 1 mL 2.5 M glycine, and crosslinked spheroplasts were pelleted, washed in cold PBS, and pelleted again prior to flash freezing. Log cells were prepared as above, except six 100 mL cultures were grown to optical density of 0.55/mL. Cells were spheroplasted in six conical tubes each of cells in 10 mL Buffer Z using 250 μl 10 mg/mL 20T zymolyase at 30°C with rotation for 30 minutes. For Micro-C XL in the absence of DSG, cells were prepared as above except they were flash frozen after formaldehyde crosslinking and again following spheroplasting. For all experiments, two conical tubes of frozen prepared spheroplasts were split into eight MNase titration reactions to determine the concentration giving approximately 95% mononucleosome-sized fragments in the insoluble chromatin fraction, and this concentration of MNase was then used to carry out MNase digestion of the remaining sample. For all experiments, MNase digestion, end repair and labeling, proximity ligation, di-nucleosomal DNA purification, and library preparation were then carried out in 10 reactions for quiescent cells and 4 reactions for log cells exactly as described in Hsieh, *et al*., 2016 (*5*), except that during library preparation, adapter ligation was completed overnight at room temperature. We found that this small modification increases the yield of unique di-nucleosome fragments in our hands. Purified di-nucleosomal DNA following the gel extraction step was combined from all 10 reactions for quiescent cells and all 4 for log prior to the library amplification. Micro-C XL samples were completed in two biological replicates at different times, agreement between replicates was determined by HiCRep (see below and **Fig. S1**), and then replicate sequencing data were merged. Experiments were completed to obtain a minimum of 80 million unique paired reads per merged sample following removal of rDNA and read-through di-nucleosome reads.

### ChIP-seq

#### Chromatin preparation

For log cells, approximately 70 optical density units (at A_660nm_) of cells in 100 mL rich media were crosslinked with 3 mL 37% formaldehyde for 20 minutes with rotation at room temperature, formaldehyde was attenuated with 3.3 mL 2.5 M glycine for 5 minutes, and cells were washed in cold TBS and flash frozen. For quiescent cells, approximately 300 optical density units of purified cells were resuspended in 25 mL water and crosslinked with 750 μl 37% formaldehyde as above, then attenuated using 1.25 mL glycine. Pellets were resuspended in 300 ul ice-cold Breaking Buffer (100 mM Tris pH 8, 20% glycerol, 1mM PMSF), approximately 600 μl of acid-washed glass beads were added, and cells were bead beat for 5 minutes (log cells) or 10 minutes (quiescent cells) until greater than 95% of cells were visibly broken under the microscope. Lysates were separated from beads, centrifuged at maximum speed for 1 minute, then pellet was resuspended in 1 mL ice-cold FA Buffer (50 mM HEPES-KOH pH. 7.6, 150 mM NaCl, 1 mM EDTA, 1% TritonX-100, 0.1% sodium deoxycholate, 1% PMSF). Chromatin was sonicated until fragmented to about 200 base pairs, then centrifuged twice at 16,000 g for 10 min at 4°C to remove residual insoluble material, and flash frozen and stored at −80°C. For quantification, an aliquot of chromatin was resuspended in an equal volume of Stop Buffer (20 mM Tris pH 8, 100 mM NaCl, 20 mM EDTA, 1% SDS) and incubated at 65°C overnight to remove crosslinks. Chromatin was then treated with 0.2 mg/mL RNaseA for 1 hour at 42°C and 0.2 mg/mL Proteinase K for 4 hours at 55°C, purified using the Qiagen MinElute PCR Cleanup Kit (catalog #28004), and quantified using a Qubit system.

#### ChIP-seq

Antibodies were conjugated to 20 μl per sample of magnetic beads with shaking for 1 hour at 20°C. For H3 ChIP, 1 μL of anti-H3 antibody (Abcam, catalog #1791) was conjugated to Dynabeads Protein G beads (Invitrogen, catalog #10004D) per reaction. For Pol II ChIP, 2 μL anti-Rpb3 antibody (Biolegend, catalog #665003) was conjugated to Dynabeads M-280 sheep anti-mouse IgG beads (Invitrogen, catalog #11201D). For FLAG ChIP-seq (for Brn1 and Hho1), 4 μL of anti-FLAG antibody (Sigma #F1804) was conjugated to Protein G beads, and for penta-H4 acetylation ChIP, 2 μl anti-penta-H4Ac antibody (Sigma, #06-946) was conjugated to Protein G beads. Beads were washed in PBST, then added to 1 μg of chromatin. Chromatin was incubated with beads with rotation at room temperature for 1.5 hours, then beads were washed three times in FA Buffer, 3 times in FA-High Salt Buffer (50 mM HEPES-KOH pH. 7.6, 500 mM NaCl, 1 mM EDTA, 1% TritonX-100, 0.1% sodium deoxycholate), and 1 time in RIPA Buffer (10 mM Tris pH 8, 0.25 M LiCl, 0.5% IGEPAL, 0.5% sodium deoxycholate, 1 mM EDTA). To elute, 50 μl of Stop Buffer was added and beads were incubated at 75°C for 15 minutes, twice. Elutions were combined and crosslinks were reversed overnight at 65°C. Chromatin was treated with RNase and PK and cleaned up as described. ChIP-seq libraries were prepared using the Ovation Ultralow v2 kit (Tecan, catalog #0344). Single-end sequencing was completed on an Illumina HiSeq 2500 in rapid run mode.

### Antibodies used in ChIP-seq and Western blot

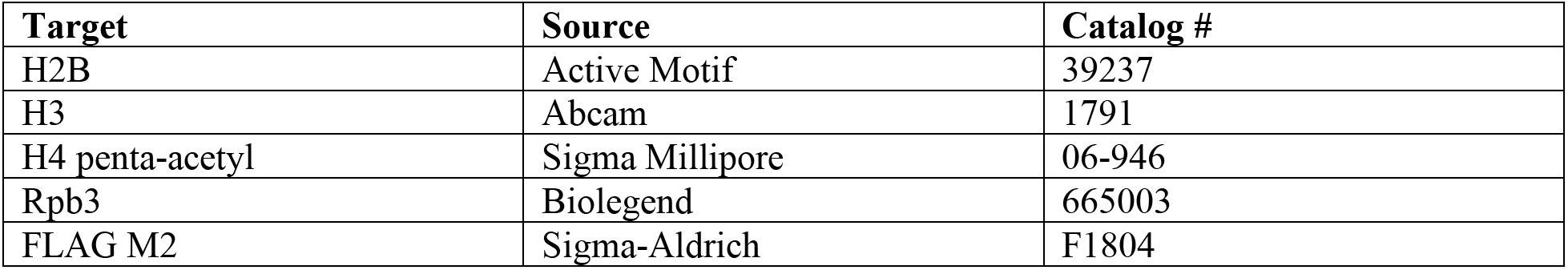

### DAPI staining and confocal microscopy

For DAPI staining, 5 optical density units at A_660nm_ of cells were fixed in 1 mL of 3.7% formaldehyde in 0.1 M KPO4 pH 6.4 at 4 C for 20 minutes with rotation. Cells were washed once in 0.1 M KPO4 pH 6.4, re-suspended in 1 mL of sorbitol/citrate (1.2 M sorbitol, 100 mM K2HPO4, 36.4 mM citric acid). Cells were then digested at 30°C in 200 μL of sorbitol/citrate containing 0.25 μg (for G1) or 2.5 μg (for quiescent) of 100T zymolyase for 5 minutes (log) or 30 minutes (quiescent). Cells were washed once and resuspended in sorbitol/citrate, then loaded onto PTFE printed slides from Electron Microscopy Sciences (catalog #63430-04) coated with 0.1% polylysine. Slides were incubated in ice cold methanol for 3 minutes and ice-cold acetone for 10 seconds, before air-drying. Then 10 μL of DAPI-mount (0.1 μg/mL DAPI, 9.25 mM p-phenylenediamine, dissolved in PBS and 90% glycerol) was added to each slide well. Z stack images with a 0.1 micron interval were obtained using a Leica TCS SP8 confocal microscope at 630X and optimized resolution. Nuclear volumes were measured using the 3D Objects Counter tool in Fiji (*46*). Threshold was set to best cover the nuclear chromatin of a maximum intensity Z-Projection. Outliers were defined as any data points less than the lower bound (Q1 − 1.5 IQR) or greater than the upper bound (Q3 + 1.5 IQR.)

### Quiescent cell longevity assay

Purified quiescent cells from 1-week-old cultures were inoculated into water at an optical density (A_660nm_) of 0.1 and incubated at 30°C with constant rotation. Samples were diluted into distilled water before plating onto YPD plates in triplicate. Plates were incubated at 30°C for 2 days before colony counting. Survival was determined by colony forming units (**CFU**). CFU at the first week was set to be the initial survival (100%).

### STEM sample preparation and imaging

Chicken erythrocytes were prepared as described previously (*17*). Briefly, erythrocytes were washed in wash buffer (130 mM NaCl, 5 mM KCl, and 10 mM HEPES, pH 7.3) and centrifuged into pelleted cells. Nuclei were prepared by resuspension of pelleted cells in lysis buffer (15 mM NaCl, 60 mM KCl, and 15 mM HEPES, pH 7.3, 0.1% NP-40, and 1μM PMSF) for 5 min and centrifuged at 1000 g for 3 min, resuspension and wash was repeated three times. Nuclei in lysis buffer were then treated with 2 mM MgCl2 and fixed with 2.5 % glutaraldehyde and 4% paraformaldehyde at 4℃, postfixed in 1% OsO4, dehydrated in series concentrations of ethanol and embedded in Spurr’s resin.

Yeast cells in 20 % BSA were cryofixed using High Pressure Freezer (HPM101, Leica) and transferred to freeze-substitution machine (AFS2, Leica). The frozen cells were freeze-substituted in the presence of 2% osmium tetroxide and 0.1% uranyl acetate in acetone at −78℃ for 48 hours, warmed up to −20℃ for 12 hours followed by 8 hours at room temperature. The fixative was washed out with acetone and embedded in Spurr’s resin at room temperature (*47*). Resin embedded cells were sectioned into either 90 nm sections for TEM/STEM imaging or 200 nm sections for STEM tomogram acquisition and mounted on copper grids. Sections were double stained with uranyl acetate and lead citrate at room temperature for enhancing contrast before EM acquisition. TEM and STEM imaging were performed using a FEI Tecnai microscope operated at 120 kV with a 2K by 2K CCD camera (US1000, Gatan) and a high-angle annular dark field (HAADF) detector (Model 3000, Fischione), respectively.

For STEM tomography, the 200 nm sections were immersed in a solution of 10 nm colloidal gold particles served as fiducial markers with 1% BSA for 30 seconds and air dried. To reduce missing wedge effect, the grids were loaded in a rotation sample holder (Model 2040, Fischione) for tomogram acquisition and the dual axial tilt images were collected from −60° to 60° at 2° increments. The STEM tomograms were reconstructed by simultaneous iterative reconstruction technique (SIRT) with 10 iterations implemented in IMOD software (*48*). Image segmentation processing was performed in Fiji, including contrast enhancement, image binary, and noise remove. The segmented EM sub-tomogram volumes (270 nm × 270 nm × 180 nm) were imported into Amira software (Thermo Scientific) to calculate chromatin fiber diameter using the surface thickness function (*2*).

### Mesoscale modeling

#### Mesoscale model

Chromatin fibers typical of Log and quiescent cells were modeled using our nucleosome resolution mesoscale model (*12*, *18*, *49*, *50*). Our chromatin mesoscale model combines four coarse-grained elements: the nucleosome core particle, with electrostatic charges derived by the Discrete Surface Charge Optimization algorithm (DiSCO) (*51*); linker DNA, modeled with a combined worm-like chain and bead model (*52*); flexible histone tails, coarse-grained as 5 residues per bead (*53*); and linker histones (**LH**s) H1e and H1c, with coarse-grained beads for the globular domain (6 beads) and for the C-terminal domain (22 beads for H1e and 21 for H1c) (*54*, *55*). Acetylated tails are modeled following our multiscale study on histone acetylation (*56*). There, we showed that acetylated tails are more rigid and folded than wild type tails, and that the chromatin unfolding upon acetylation occurs due to the impairment of internucleosome interactions caused by the folded and rigid tails. Thus, we use the configuration of folded tails and ensure rigidity by increasing the force constants in the energy terms by a factor of 100. Mg^2+^ presence is modeled by a phenomenological approach (*9*), in which the DNA persistence length is reduced from 50 to 30 nm based on experimental data (*57*) and the electrostatic repulsion among linker DNAs is reduced by increasing the inverse Debye length in the DNA-DNA electrostatic term.

Coarse-grained elements have bonded interactions, which consist of stretching, bending, and twisting terms. Nonbonded interactions among coarse-grained elements are modeled with the Debye-Hückel approximation to treat the electrostatics and with Lennard-Jones potentials to treat excluded volume terms. For details on model parameters and energy terms, please see (*49*).

#### Chromatin systems

To study quiescent and Log cell chromatin, we model a segment of 40 kb located between 130,000 bp and 170,000 bp of Chr1. This segment was selected based on Micro-C contact maps that show significant chromatin reorganization in this region, although the differences between Log and quiescence persist throughout the genome. Moreover, this region does not show a strong presence of condensin binding (*3*). Nucleosome positions were obtained from MNase-seq data (*25*) using the DANPOS algorithm (*58*). Nucleosomes called by DANPOS with summit values below 1% of the average summit value per condition were removed. Linker histone and histone acetylation positions were obtained from ChIP-seq data using using the “callpeak” function of the MACS2 algorithm (*59*). As a result, the Log chromatin fiber contains 222 nucleosomes, 61 nucleosomes acetylated, and a LH density of 0.05 LH/nucleosome. The quiescent chromatin fiber contains 228 nucleosomes, 3 nucleosomes acetylated, and a LH density of 0.29 LH/nucleosome (**Fig. S4A**).

#### Monte Carlo Sampling

Fibers typical of Log and quiescent cells are subject to Monte Carlo (MC) simulations starting from an ideal zigzag geometry as we have shown this configuration to be dominant under physiological salt conditions (*9*). 30 independent trajectories are run for each system for at least 60 million MC steps. Each simulation is initiated from a different random seed number and a randomly chosen B twist value for the DNA of −12°, 0°, or +12° to mimic natural variations (*60*). To mimic physiological conditions, simulations are performed in presence of 150 mM NaCl and 1mM Mg2+, and a temperature of 293 K.

Five types of tailored MC moves are implemented for the efficient global and local sampling of the fibers. A global pivot move chooses a random position along the fiber and then rotates the shorter section of the bisected chain around a randomly chosen axis running through that point. The resulting configuration is accepted or rejected based on the Metropolis criteria (*61*). All DNA and LH beads are subject to translation and rotation moves also accepted or rejected based on the Metropolis criteria. A configurationally biased regrow routine is used to simulate the rapid movement of histone tails. A randomly chosen histone tail is regrown starting with the bead closest to the core; the new configuration is accepted or rejected based on the Rosenbluth criteria (*62*). Acetylated tails are sampled with a fold-swap move in which tails are randomly chosen and its fold state is swapped. Thus, if a chosen tail is currently folded (acetylated), its coordinates and equilibrium values are swapped with those of the unfolded version of that tail (wild type). The new configuration is accepted or rejected based on the Metropolis criteria. Folded tails do not interact with any other chromatin element and are not subject to the regrow routine.

During the MC simulation, convergence of the systems is carefully checked by monitoring global and local properties (**Fig. S4B-C**). The last 10 million MC steps of each independent trajectory, corresponding to a total of 3,000 configurations, are used for analysis.

#### Analysis

The sedimentation coefficient (S_w,20_), in units of Svedbergs, is used to describe the compaction of the fiber. It is defined by the expression:

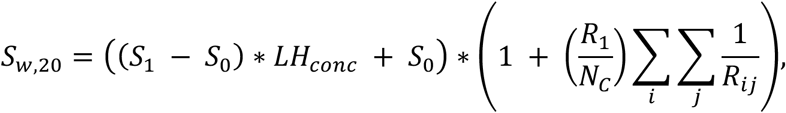

where S_0_ is the sedimentation coefficient for a mononucleosome with LH bound (12 S) (*63*), S_1_ is the sedimentation coefficient for a mononucleosome without LH (11.1 S) (*23*), LH_conc_ the concentration of LH in the fiber, R_1_ the spherical radius of a nucleosome (5.5 nm), N_C_ the number of nucleosomes in the fiber, and R_ij_ the distance between two nucleosomes i and j.

The radius of gyration, which describes the overall dimension of the polymer chain, is measured as the root mean squared distance of each nucleosome from the center of mass according to the relation:

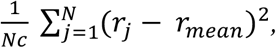

where N_c_ is the number of nucleosomes, r_j_ the center position of the nucleosome core j, and r_mean_ the average of all core positions (*50*).

Fiber volumes are calculated using the AlphaShape function of Matlab, which creates a nonconvex bounding volume that envelops the nucleosomes. Surfaces are visually inspected to ensure that they represent correctly the fiber morphology (**Fig. S4D**). This is because non cylindrical-like shapes may not well be estimated. In that case, the AlphaShape object can be manipulated to tighten or loosen the fit around the points to create a nonconvex region.

Packing ratio is used to describe the compaction of the fiber and is measured as the number of nucleosomes contained in 11 nm of fiber. It is determined according to the relation:

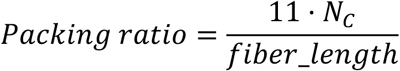

where N_C_ is the number of nucleosomes and the fiber length is calculated using a cubic smoothing spline function native from Matlab.

#### Internucleosome interactions

Internucleosome contacts were calculated and reported every 10,000 steps during the simulation. A contact is defined if the tails or charge beads of nucleosome i are found to be within 2 nm of the tails or charge beads of nucleosome j. Contact maps for each trajectory are normalized by the maximal number of contacts seen throughout the trajectory, and the resulting normalized frequencies are summed together.

These internucleosome matrices are projected into normalized one-dimensional plots that depict the relative intensity of interactions between nucleosomes separated by *k* neighbors as follows:

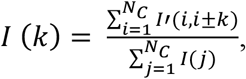

where Nc is the number of nucleosomes and I’ the internucleosome interaction matrix.

#### Tail interactions

The interactions of tails with other chromatin components such as DNA, nucleosomes, and other tails, are calculated similarly as described above. For each trajectory, we measure the fraction of the MC steps that each tail (H2A, H2B, H3, and H4) is “in contact” with non-parental nucleosome, non-parental DNA, or other tails of a non-parental nucleosome. Contacts are defined when the distance between the beads of both elements is less than 2 nm. The interaction frequencies are averaged over the 30 trajectories for each system, obtaining means and standard deviations.

### Micro-C XL data analysis

#### Sequencing read processing

The two ends of paired-end Micro-C reads were mapped independently to the sacCer3 reference genome (release R64-2-1) using bowtie2 version 2.3.5.1 with the “--very-sensitive” parameter set (*64*). All read pairs where either end received MAPQ score < 6 were removed. All the remaining in-facing read pairs were removed. The resulting read pairs were processed into the multiple-bin size contact matrix in the Cooler format (https://github.com/mirnylab/cooler). The bin sizes we used in the downstream analyses were 10 bp, 200 bp and 5000 bp. Micro-C heatmaps were generated using Juicebox (*65*).

#### Contact probability decay curve

Each diagonal of the 10-bp Cooler contact matrix contains the Micro-C interactions between genomic loci at the genomic distance of a multiple of 10 bp. The contact counts in each diagonal were summed and the sum was then divided by the number of elements in the diagonal. The result is then normalized so that the contact probability decay sums to one across all genomic distances. Analysis was completed separately for each orientation of ligation pairs (“in,” “out,” and “same,” which includes “in-out” and “out-in” pairs). “Same” contacts are shown in all figures as these showed the most consistent nucleosome phasing across samples.

#### Micro-C contacts pileup analysis

We used the Micro-C contact matrix of 200-bp bin size to perform pileup analysis. Previous (*3*) work defined the condensin ChIP peaks and L-CID boundaries [L-CID boundaries were previously called using the cworld-dekker package using matrix2insulation.pl with settings– is4800–nt0.4–ids3200–ss800–im mean (*66*)]. A 20 kb window centering on each of the condensin ChIP peaks/L-CID boundaries is defined as a target region in this analysis. The set of Micro-C contacts between two target regions is defined as the submatrix of contacts between the two target regions. We call it an inter-peak submatrix if the two target regions correspond to two different condensin ChIP peaks/L-CID boundaries, or an intra-peak submatrix if the two target regions correspond to the same condensin ChIP peak/L-CID boundary. The element-wise median across a set of submatrices is defined as the pileup matrix. We call it an inter-peak pileup matrix if all the submatrices involved are inter-peak submatrices, or an intra-peak pileup matrix if all the submatrices are intra-peak submatrices.

#### HiCRep analysis

We implemented the HiCRep algorithm (*15*) in Python (*16*) to compute the stratum-adjusted correlation coefficient (SCC) between two Micro-C contact matrices. The 5000-bp bin size contact matrices are used in this analysis. The input contact matrix is first normalized by dividing the contact counts by the sum of all contacts in the matrix. Then the chromosome-wise SCC scores between two normalized matrices are computed using the contacts up to 100 kb of genomic distance. The median of the chromosome SCC scores between the two matrices is reported.

### ChIP-seq data analysis

Reads were aligned to the sacCer3 reference genome (release R64-2-1) using bowtie2 in “--very-sensitive” mode (*64*), then filtered and indexed using SAMtools (*67*). Bam files were then RPKM normalized and converted to bigwig files using the “bamCompare” command in deepTools2.0 (*68*), and IPs were normalized to inputs using “bigwigCompare.” Heatmaps, metaplots, and Pearson correlations were also generated using deepTools. Genome browser views were generated using the Integrated Genome Browser software (*69*). Peak calling was completed using “callpeak” and “bdgdiff” commands in MACS2 (*59*). For final analysis, fastq files from two biological replicates were merged for each condition.

---

**Fig. S1.**
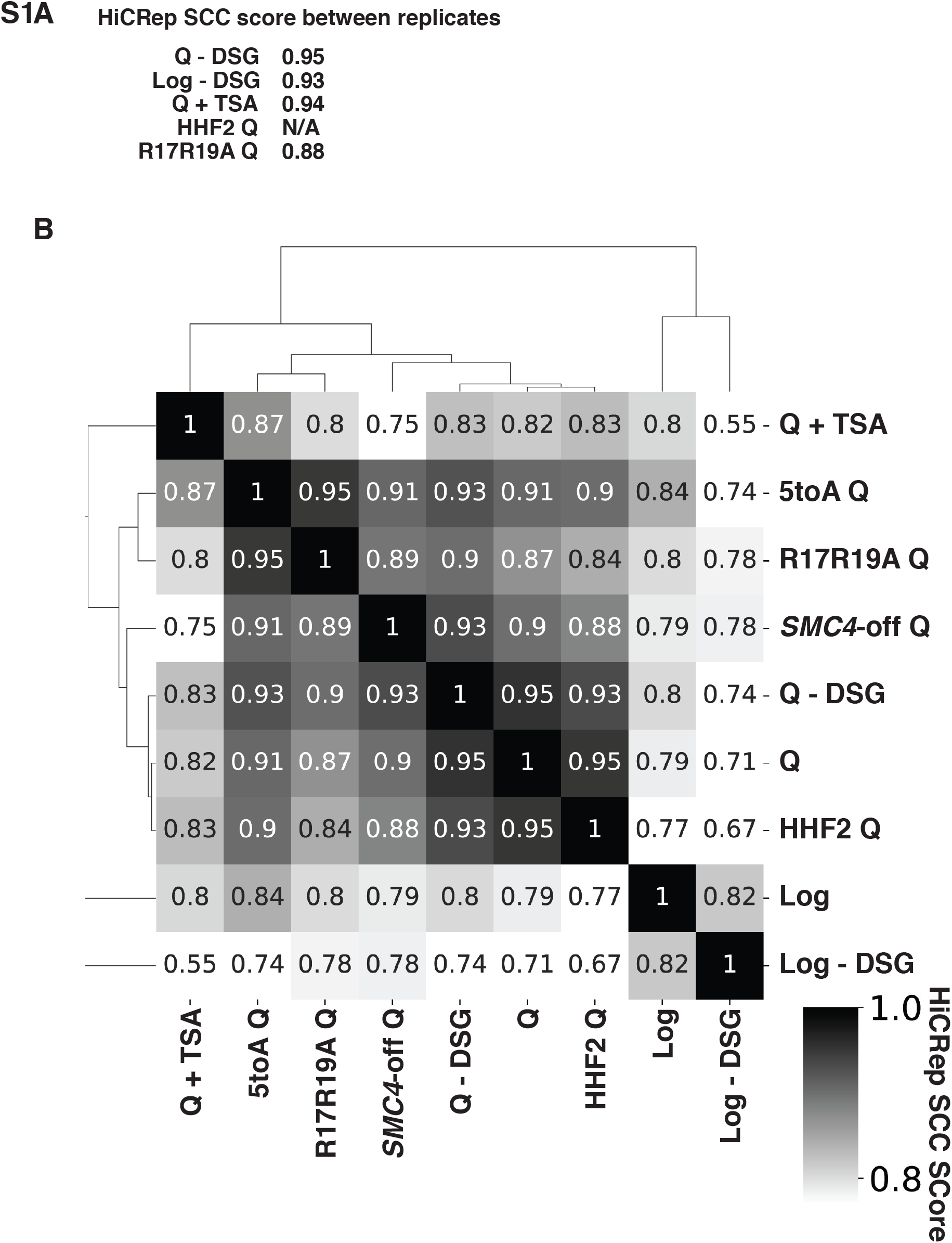
HiCRep scores of Micro-C data. (**A**) Stratum-adjusted correlation coefficients (SCC) calculated by HiCRep of Micro-C XL data from two biological replicates of each of the indicated conditions. HHF2 was completed in only one replicate to determine similarity to true WT. (**B**) SCCs calculated and hierarchically sorted between conditions. Micro-C XL data of biological replicates were merged prior to analysis. Log, quiescent (Q), and *SMC4*-off cell Micro-C XL data were previously published in (*3*). Log and Q represent data from true WT strains.

**Fig. S2.**
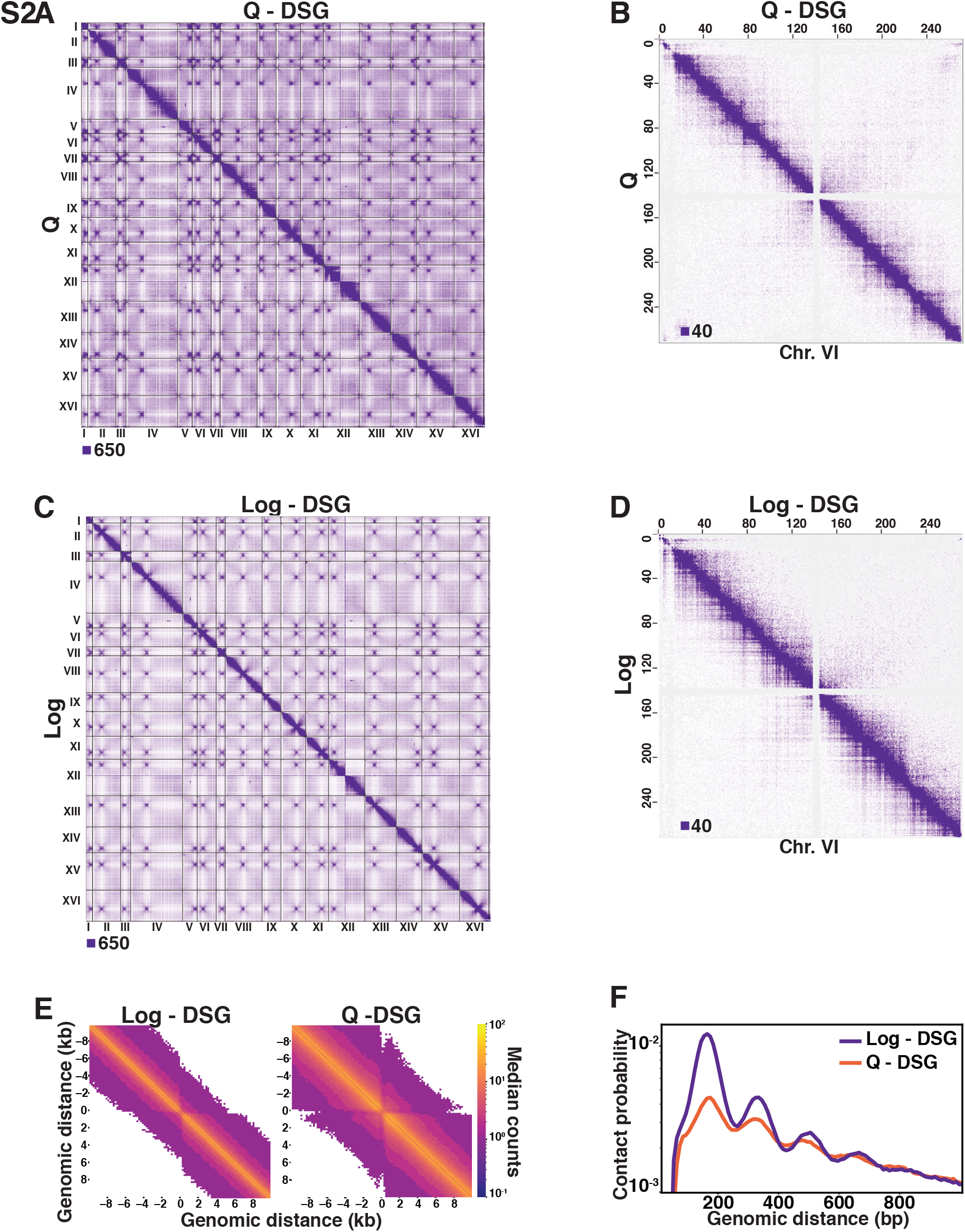
Omitting DSG from the Micro-C XL protocol diminishes contacts in log cells. (**A**) Genome-wide Micro-C XL data in Q with and without DSG. (**B**) Micro-C XL data at 1 kb resolution. (**C**) Genome-wide Micro-C XL data in Log with and without DSG. (**D**) Micro-C XL data at 1 kb resolution. (**E**) Micro-C XL metaplots of median interactions +/−10 kb around sites of condensin-bound L-CID boundaries at 200 bp resolution. The scale shows median counts. (**F**) Contact probability map of Micro-C XL short crosslinker data in Log and Q.

**Fig. S3.**
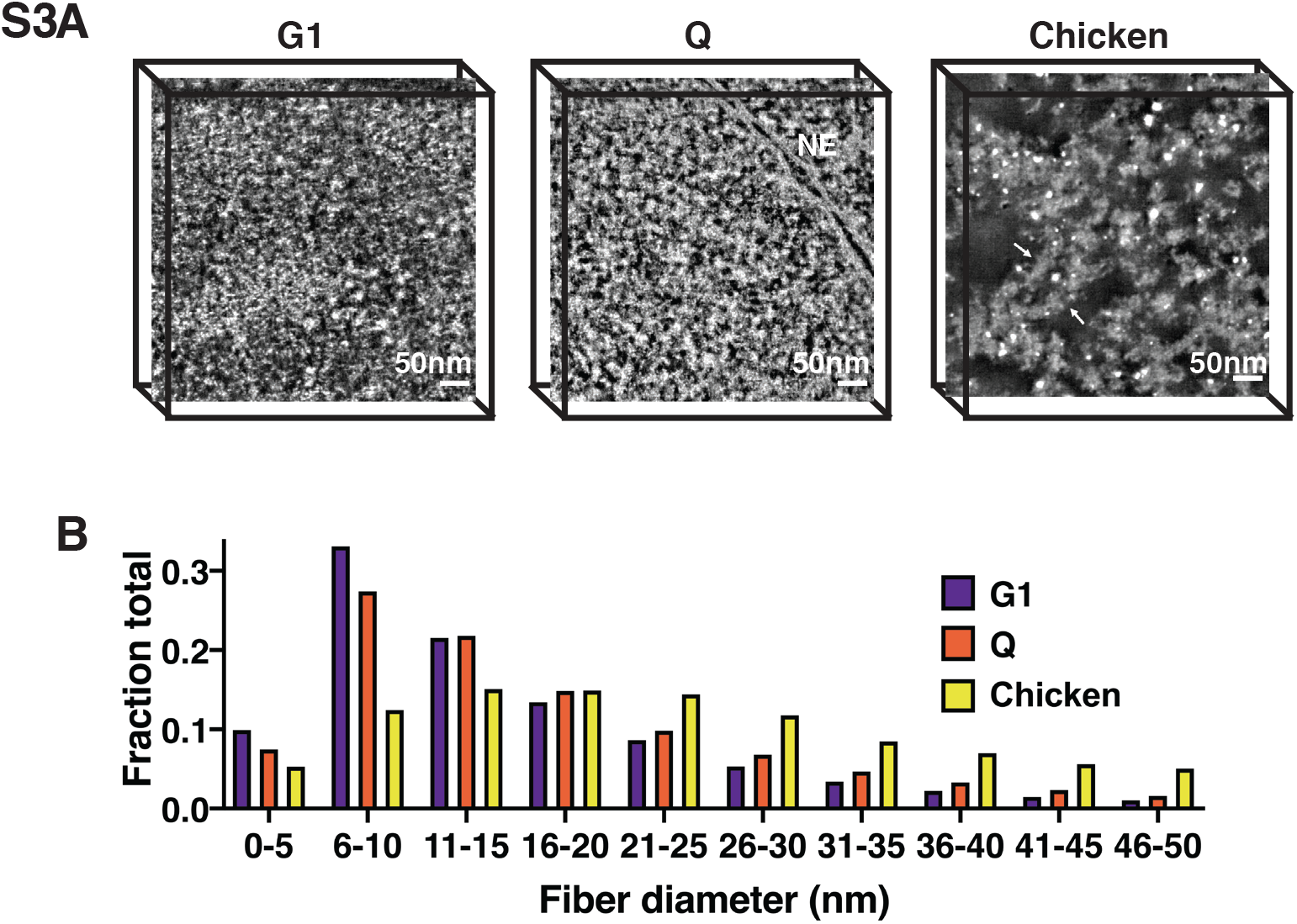
STEM tomography of G1 and Q yeast and chicken erythrocyte nuclei. (**A**) HAADF-STEM images of uranyl acetate and lead citrate stained G1-arrested, Q cell, and magnesium-treated chicken erythrocyte nuclei slices. Chromatin fibers appear as white. NE is nuclear envelope. Arrows point to 30 nm fibers. Scale bar represents 50 nm. (**B**) Histograms of fiber diameter counts calculated using the surface thickness function in Amira software.

**Fig. S4.**
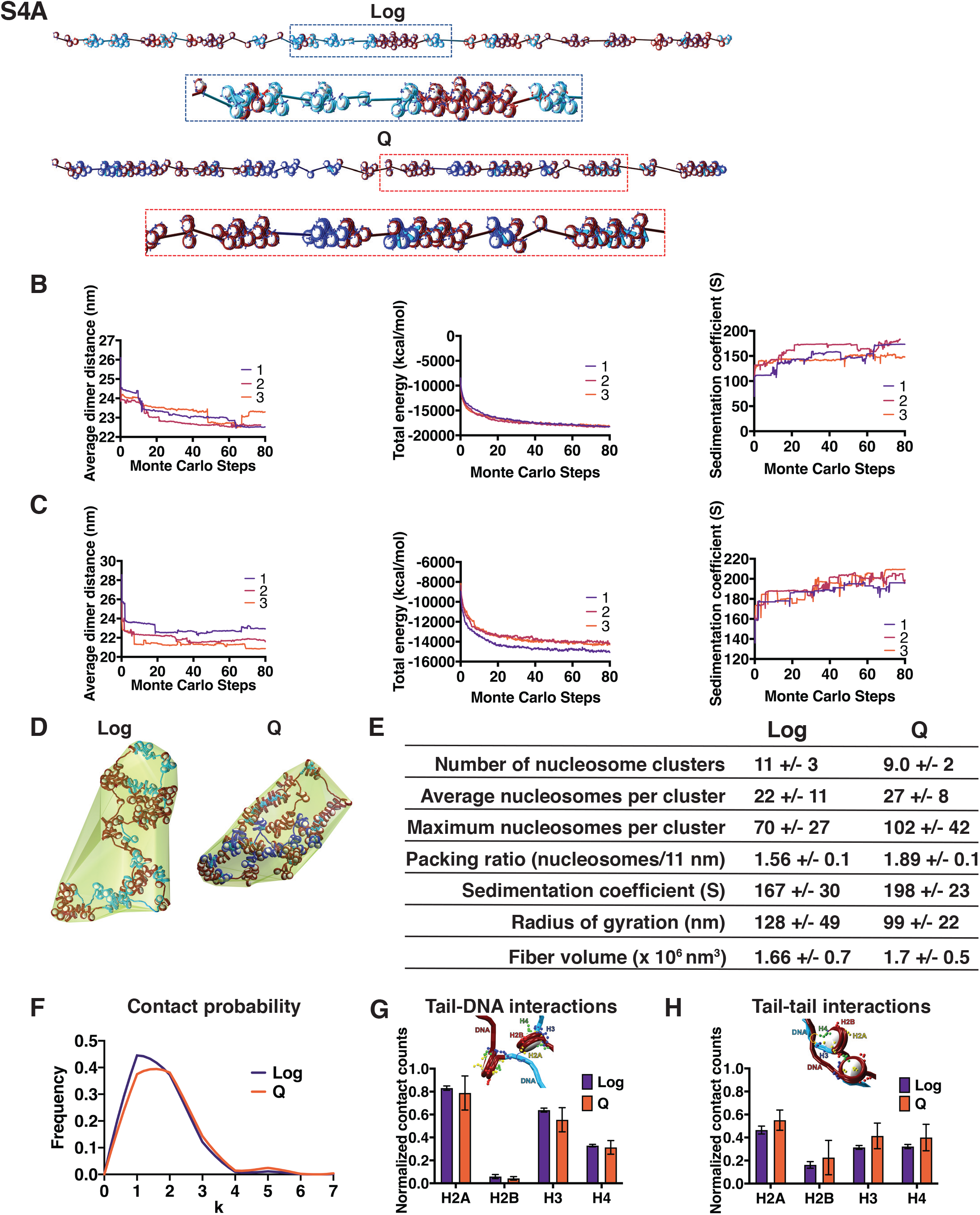
Mesoscale modeling of Log and Q chromatin fibers. (**A**) Starting configurations for Q and Log cell chromatin fibers. Genes are shown in light blue and blue for Log and Q, respectively. For both systems, intergenic regions are shown in dark red, linker histone in turquoise, wild type histone tails in blue, and acetylated histone tails in red. (**B-C**) Simulation convergence assessed through global and local properties. Shown is the average dimer distance, total energy, and sedimentation coefficient along 80 million MC steps. We show the convergence for three independent trajectories for log (**B**) and Q (**C**). (**D**) Representative final configurations showing bounding surface. Genes are shown in blue and intergenic regions are shown in red. Hho1 is shown in turquoise. (**E**) Fiber compaction and morphology parameters for the 3000-configuration ensemble of each system. (**F**) Modeled frequency of nucleosome contact probabilities between a nucleosome and each subsequent nucleosome (k). (**G**) Normalized counts of histone tail/non-parental DNA contacts averaged across 30 independent trajectories. Error bars show standard deviation. (**H**) Normalized counts of histone tail/histone tail contacts averaged across 30 independent trajectories. Error bars show standard deviation.

**Fig. S5.**
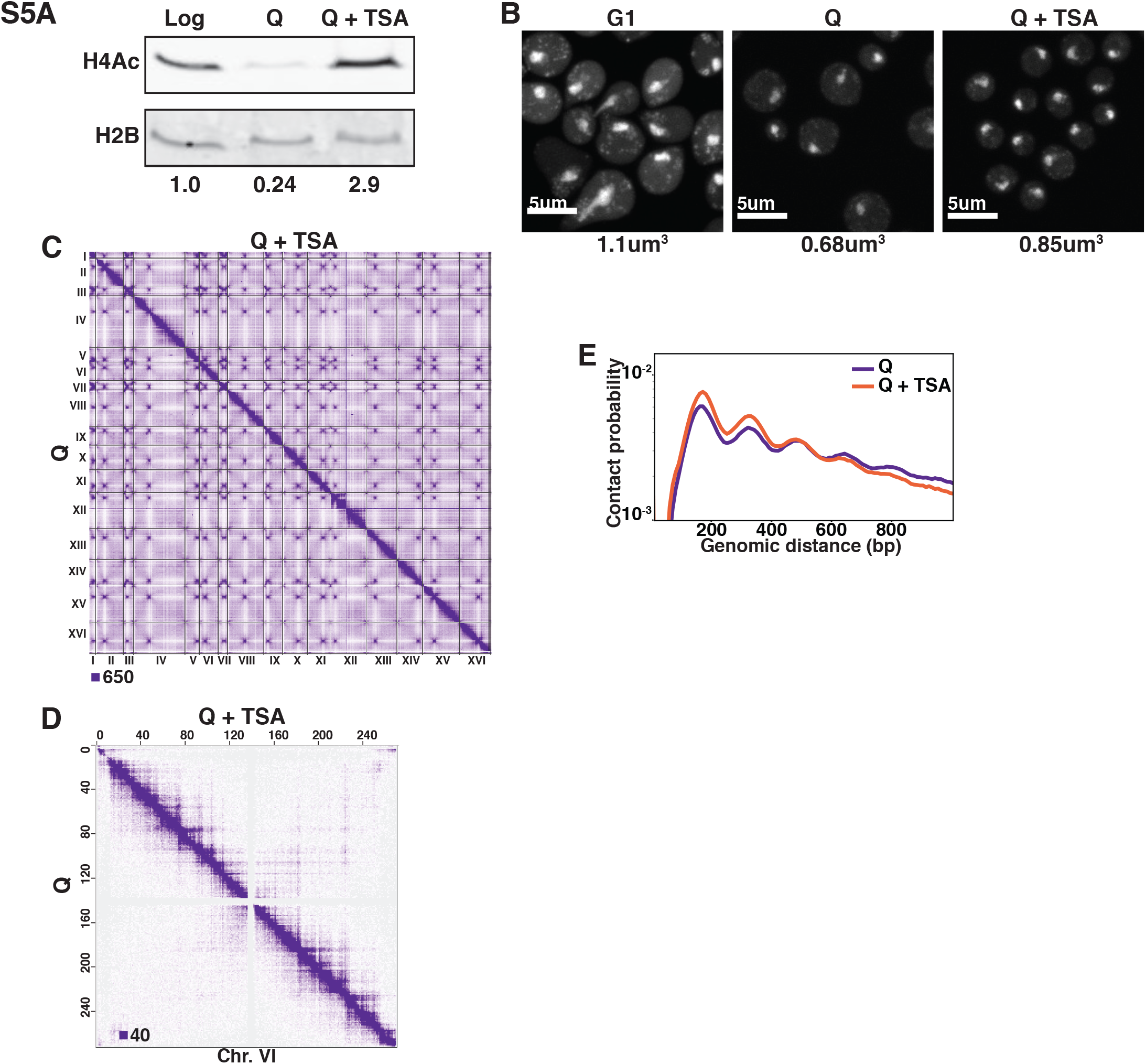
TSA treatment increases H4 tail acetylation and decompacts chromatin in Q cells. (**A**) Representative Western blots of H4 penta-acetylation (H4Ac) and H2B in Log, Q, and TSA-treated Q cells. Band volumes were quantified using Fiji and H4Ac volumes were normalized to H2B volumes. (**B**) Representative images of DAPI-stained cells. Chromatin volumes were calculated as described in Figure 2 and in the Materials and Methods. Mean volumes are shown below. (**C**) Genome-wide Micro-C XL data in Q cells with (top) and without (bottom) TSA treatment. (**D**) Micro-C XL data at 1 kb resolution with (top) and without (bottom) TSA treatment. (**E**) Contact probability map of Micro-C XL data.

**Fig. S6.**
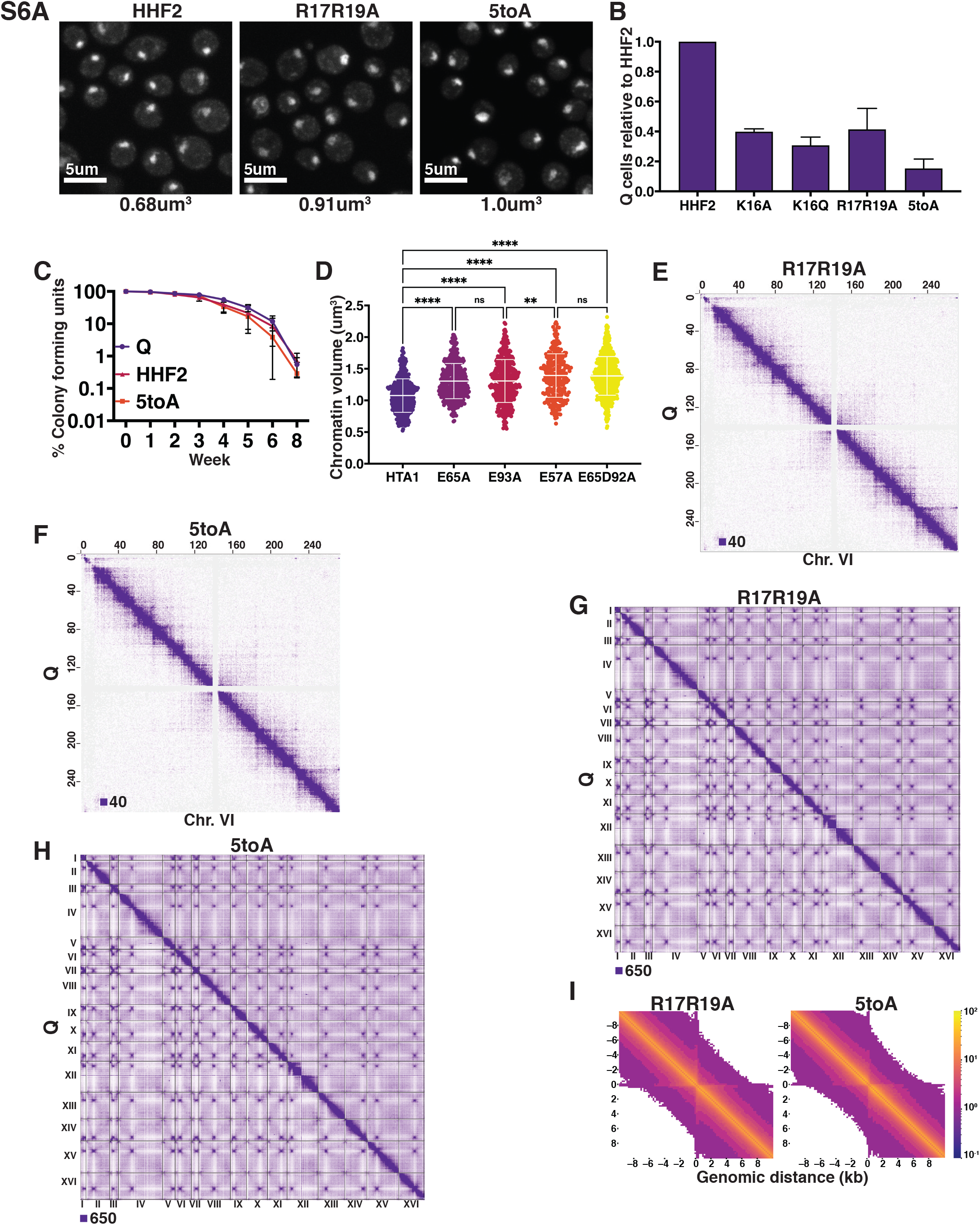
H4 basic patch and H2A acidic patch substitutions decompact chromatin in Q. (**A**) Representative images of DAPI-stained cells. Chromatin volumes were calculated as described in Figure 2 and in the Materials and Methods. Mean volumes are shown below. HHF2 is a WT control. (**B**) Normalized optical density units of purified Q cells of two biological replicates of the indicated condition to show Q entry efficiency. Error bars represent standard deviation. (**C**) The longevity of purified Q cells was measured by following the ability of cells to form colonies after the indicated number of weeks. Numbers shown represent two to three technical replicates each of two biological replicates. Error bars show standard deviation. (**D**) Chromatin volume measurements of H2A mutant Q cells following DAPI staining of at least 200 cells each of two biological replicates. Bars represent mean and standard deviation. Significance was determined using two-tailed paired t-tests. HTA1 is WT. Note: As H2A mutants were generated in auxotrophic strain backgrounds in which chromatin volumes are greater than in prototrophic backgrounds, these data cannot be directly compared to other DAPI datasets in this manuscript. (**E** and **F**) Genome-wide Micro-C XL data. (**G** and **H**) Micro-C XL data at 1 kb resolution. (**I**) Micro-C XL metaplots of median interactions +/−10 kb around sites of condensin-bound L-CID boundaries at 200 bp resolution. The scale shows median counts.

**Fig. S7.**
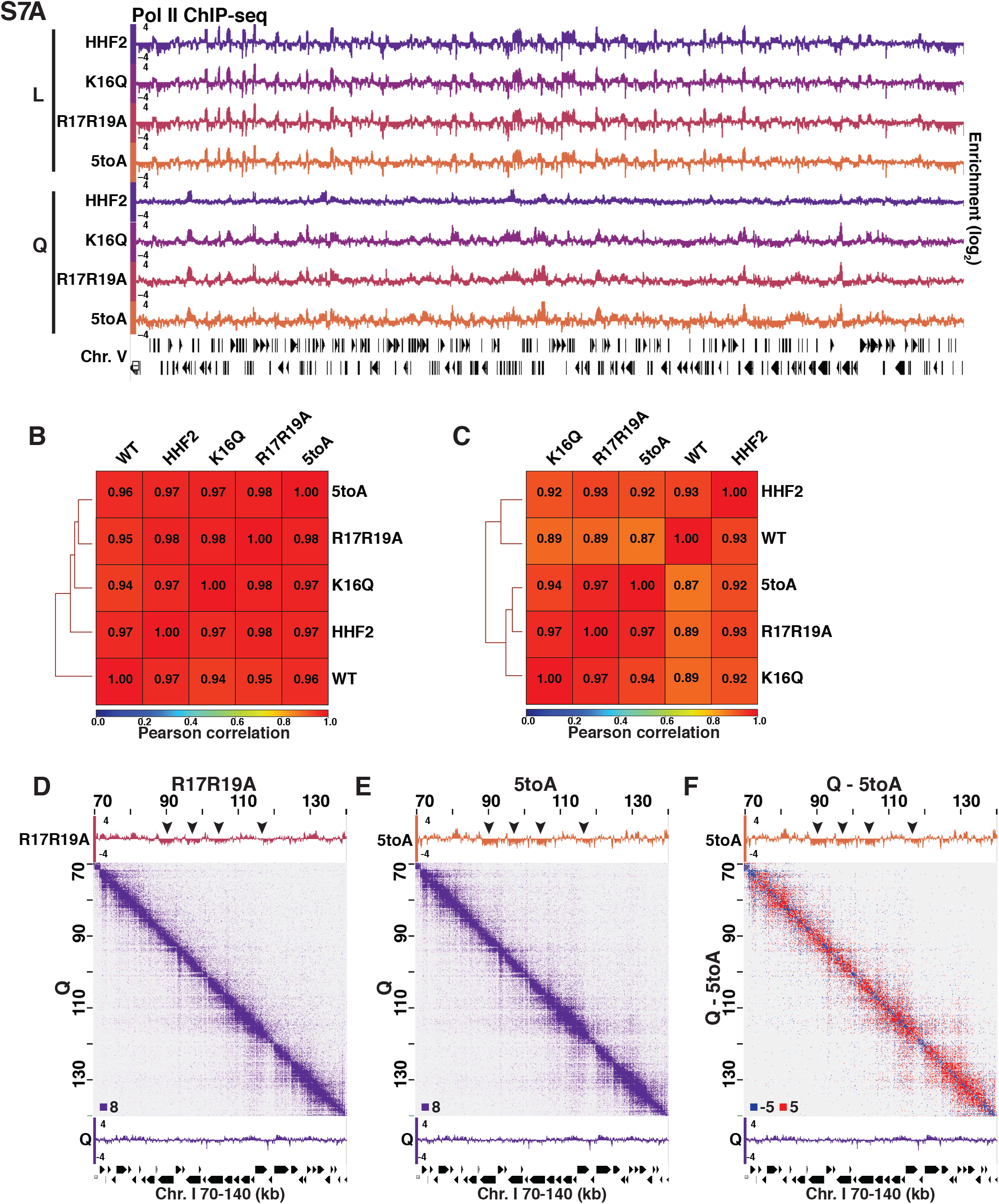
Pol II and H3 ChIP-seq in Log and Q. (**A**) Genome browser view of Pol II subunit Rpb3 ChIP-seq data in log and quiescent mutant strains across the entirety of Chromosome V. (B) Pearson correlation scores calculated and hierarchically sorted using deepTools of Rpb3 (Pol II) ChIP-seq data in log and (**C**) Q. (**D-F**) Micro-C XL and Pol II ChIP-seq data in mutant and WT quiescent (Q) cells. Arrowheads point to regions that are decompacted but not expressed in mutant cells. Micro-C XL data subtraction is shown in (**F**).

**Fig. S8.**
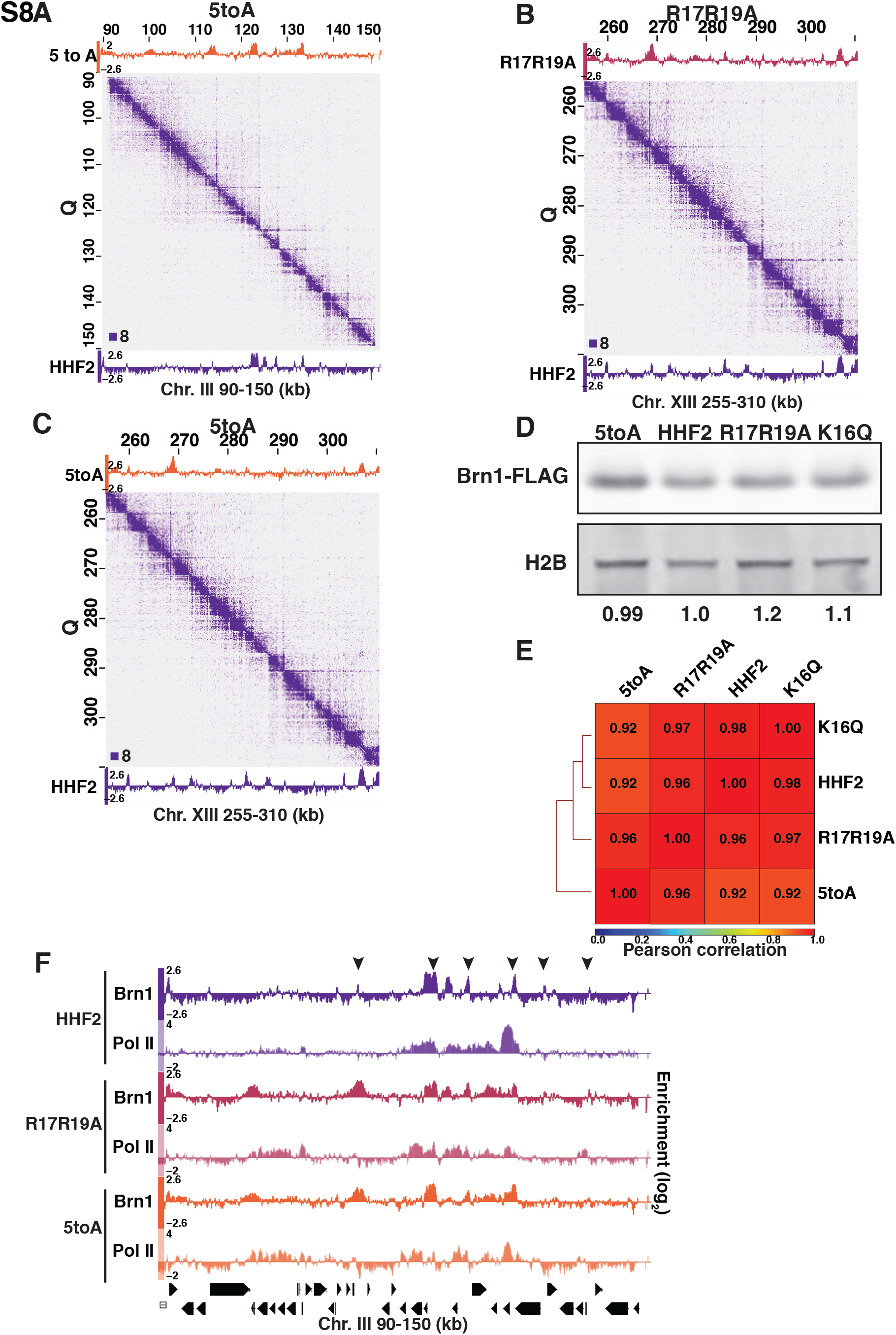
H4-mediated chromatin fiber folding represses condensin loop extrusion. (**A-C**) Condensin subunit Brn1 ChIP-seq in R17R19A (top) and HHF2 (bottom) data overlayed across Micro-C XL data at 200 bp resolution. (**D**) Representative Western blots of Brn1-FLAG and H2B in H4 mutant and HHF2 (WT) strains. Band volumes were quantified using Fiji and Brn1-FLAG volumes were normalized to H2B volumes. (**E**) Pearson correlation scores calculated and hierarchically sorted using deepTools of Brn1-FLAG in H4 mutant and HHF2 (WT) strains. (**F**) Genome browser view of Pol II subunit Rpb3 and Brn1-FLAG ChIP-seq data in H4 mutant strains across part of Chromosome III. Positions of stripes are indicated with arrowheads.

## References and Notes

1. N. Friedman, O. J. Rando, Epigenomics and the structure of the living genome. Genome Res. 25, 1482–1490 (2015).

2. H. D. Ou, S. Phan, T. J. Deerinck, A. Thor, M. H. Ellisman, C. C. O’Shea, ChromEMT: Visualizing 3D chromatin structure and compaction in interphase and mitotic cells. Science 357, eaag0025 (2017), doi:10.1126/science.aag0025.

3. S. G. Swygert, S. Kim, X. Wu, T. Fu, T.-H. Hsieh, O. J. Rando, R. N. Eisenman, J. Shendure, J. N. McKnight, T. Tsukiyama, Condensin-Dependent Chromatin Compaction Represses Transcription Globally during Quiescence. Mol. Cell. 73(2019), doi:10.1016/j.molcel.2018.11.020.

4. T. H. S. Hsieh, A. Weiner, B. Lajoie, J. Dekker, N. Friedman, O. J. Rando, Mapping Nucleosome Resolution Chromosome Folding in Yeast by Micro-C. Cell. 162, 108–119 (2015).

5. T. H. S. Hsieh, G. Fudenberg, A. Goloborodko, O. J. Rando, Micro-C XL: Assaying chromosome conformation from the nucleosome to the entire genome. Nat. Methods. 13, 1009–1011 (2016).

6. C. Allen, S. Büttner, A. D. Aragon, J. A. Thomas, O. Meirelles, J. E. Jaetao, D. Benn, S. W. Ruby, M. Veenhuis, F. Madeo, M. Werner-Washburne, Isolation of quiescent and nonquiescent cells from yeast stationary-phase cultures. J. Cell Biol. 174, 89–100 (2006).

7. I. Sagot, D. Laporte, The cell biology of quiescent yeast – a diversity of individual scenarios. J. Cell Sci. 132(2019), doi:10.1242/jcs.213025.

8. D. Lohr, G. Ide, Comparison of the structure and transcriptional capability of growing phase and stationary yeast chromatin: A model for reversible gene activation. Nucleic Acids Res. 6, 1909–1927 (1979).

9. S. A. Grigoryev, G. Arya, S. Correll, C. L. Woodcock, T. Schlick, Evidence for heteromorphic chromatin fibers from analysis of nucleosome interactions. Proc. Natl. Acad. Sci. U. S. A. 106, 13317–13322 (2009).

10. B. Dorigo, T. Schalch, A. Kulangara, S. Duda, R. R. Schroeder, T. J. Richmond, Nucleosome arrays reveal the two-start organization of the chromatin fiber. Science 306, 1571–1573 (2004).

11. T. Schalch, S. Duda, D. F. Sargent, T. J. Richmond, X-ray structure of a tetranucleosome and its implications for the chromatin fibre. Nature. 436, 138–141 (2005).

12. R. Collepardo-Guevara, T. Schlick, Chromatin fiber polymorphism triggered by variations of DNA linker lengths. Proc. Natl. Acad. Sci. U. S. A. 111, 8061–8066 (2014).

13. P. J. J. Robinson, L. Fairall, V. A. T. Huynh, D. Rhodes, EM measurements define the dimensions of the “30-nm” chromatin fiber: Evidence for a compact, interdigitated structure. Proc. Natl. Acad. Sci. U. S. A. 103, 6506–6511 (2006).

14. S. A. Grigoryev, G. Bascom, J. M. Buckwalter, M. B. Schubert, C. L. Woodcock, T. Schlick, Hierarchical looping of zigzag nucleosome chains in metaphase chromosomes. Proc. Natl. Acad. Sci. U. S. A. 113, 1238–1243 (2016).

15. T. Yang, F. Zhang, G. G. Yardımci, F. Song, R. C. Hardison, W. S. Noble, F. Yue, Q. Li, HiCRep: assessing the reproducibility of Hi-C data using a stratum-adjusted correlation coefficient. Genome Res. 27, 1939–1949 (2017).

16. D. Lin, J. Sanders, W. S. Noble, bioRxiv, in press, doi:10.1101/2020.10.27.357756.

17. J. Langmore, C. Schutt, The higher order structure of chicken erythrocyte chromosomes in vivo. Nature. 288, 620–622 (1980).

18. G. Arya, T. Schlick, Role of histone tails in chromatin folding revealed by a mesoscopic oligonucleosome model. Proc. Natl. Acad. Sci. U. S. A. 103, 16236–16241 (2006).

19. G. D. Bascom, C. G. Myers, T. Schlick, Mesoscale modeling reveals formation of an epigenetically driven HOXC gene hub. Proc. Natl. Acad. Sci. U. S. A. 116, 4955–4962 (2019).

20. B. Dorigo, T. Schalch, K. Bystricky, T. J. Richmond, Chromatin fiber folding: Requirement for the histone H4 N-terminal tail. J. Mol. Biol. 327, 85–96 (2003).

21. P.-Y. Kan, T. L. Caterino, J. J. Hayes, The H4 Tail Domain Participates in Intra- and Internucleosome Interactions with Protein and DNA during Folding and Oligomerization of Nucleosome Arrays. Mol. Cell. Biol. 29, 538–546 (2009).

22. P. M. Schwarz, A. Felthauser, T. M. Fletcher, J. C. Hansen, Reversible oligonucleosome self-association: Dependence on divalent cations and core histone tail domains. Biochemistry. 35, 4009–4015 (1996).

23. G.-R. M, D. F, A. J., Role of the histone “tails” in the folding of oligonucleosomes depleted of histone H1. J. Biol. Chem. 267, 19587–95 (1992).

24. M. Shogren-Knaak, H. Ishii, J. M. Sun, M. J. Pazin, J. R. Davie, C. L. Peterson, Histone H4-K16 acetylation controls chromatin structure and protein interactions. Science 311, 844–847 (2006).

25. J. N. McKnight, J. W. Boerma, L. L. Breeden, T. Tsukiyama, Global Promoter Targeting of a Conserved Lysine Deacetylase for Transcriptional Shutoff during Quiescence Entry. Mol. Cell. 59, 732–743 (2015).

26. J. Otterstrom, A. Castells-Garcia, C. Vicario, P. A. Gomez-Garcia, M. P. Cosma, M. Lakadamyali, Super-resolution microscopy reveals how histone tail acetylation affects DNA compaction within nucleosomes in vivo. Nucleic Acids Res. 47, 8470–8484 (2019).

27. M. A. Ricci, C. Manzo, M. F. García-Parajo, M. Lakadamyali, M. P. Cosma, Chromatin fibers are formed by heterogeneous groups of nucleosomes in vivo. Cell. 160, 1145–1158 (2015).

28. D. Laporte, F. Courtout, S. Tollis, I. Sagot, Quiescent Saccharomyces cerevisiae forms telomere hyperclusters at the nuclear membrane vicinity through a multifaceted mechanism involving Esc1, the Sir complex, and chromatin condensation. Mol. Biol. Cell. 27, 1875–1884 (2016).

29. S. G. Swygert, T. Tsukiyama, Unraveling quiescence-specific repressive chromatin domains. Curr. Genet. 65, 1145–1151 (2019).

30. S. Yamasaki, P. Anderson, Reprogramming mRNA translation during stress. Curr. Opin. Cell Biol. 20, 222–226 (2008).

31. L. Li, S. Miles, Z. Melville, A. Prasad, G. Bradley, L. L. Breeden, Key events during the transition from rapid growth to quiescence in budding yeast require posttranscriptional regulators. Mol. Biol. Cell. 24, 3697–3709 (2013).

32. T. G. Fazzio, M. E. Gelbart, T. Tsukiyama, Two Distinct Mechanisms of Chromatin Interaction by the Isw2 Chromatin Remodeling Complex In Vivo. Mol. Cell. Biol. 25, 9165–9174 (2005).

33. A. Goloborodko, M. V. Imakaev, J. F. Marko, L. Mirny, Compaction and segregation of sister chromatids via active loop extrusion. Elife. 5(2016), doi:10.7554/eLife.14864.

34. E. J. Banigan, A. A. van den Berg, H. B. Brandão, J. F. Marko, L. A. Mirny, Chromosome organization by one-sided and two-sided loop extrusion. Elife. 9(2020), doi:10.7554/eLife.53558.

35. L. Vian, A. Pękowska, S. S. P. Rao, K. R. Kieffer-Kwon, S. Jung, L. Baranello, S. C. Huang, L. El Khattabi, M. Dose, N. Pruett, A. L. Sanborn, A. Canela, Y. Maman, A. Oksanen, W. Resch, X. Li, B. Lee, A. L. Kovalchuk, Z. Tang, S. Nelson, M. Di Pierro, R. R. Cheng, I. Machol, B. G. St Hilaire, N. C. Durand, M. S. Shamim, E. K. Stamenova, J. N. Onuchic, Y. Ruan, A. Nussenzweig, D. Levens, E. L. Aiden, R. Casellas, The Energetics and Physiological Impact of Cohesin Extrusion. Cell. 173, 1165–1178.e20 (2018).

36. J. Nuebler, G. Fudenberg, M. Imakaev, N. Abdennur, L. A. Mirny, Chromatin organization by an interplay of loop extrusion and compartmental segregation. Proc. Natl. Acad. Sci. U. S. A. 115, E6697–E6706 (2018).

37. S. Zhang, N. Übelmesser, N. Josipovic, G. Forte, J. A. Slotman, H. Gothe, E. Gade Gusmao, C. Becker, J. Altmüller, V. Roukos, K. S. Wendt, D. Marenduzzo, A. Papantonis, bioRxiv, in press, doi:10.1101/2020.10.27.356915.

38. A. Lengronne, Y. Katou, S. Mori, S. Yokabayashi, G. P. Kelly, T. Ito, Y. Watanabe, K. Shirahige, F. Uhlmann, Cohesin relocation from sites of chromosomal loading to places of convergent transcription. Nature. 430, 573–578 (2004).

39. N. T. Tran, M. T. Laub, T. B. K. Le, SMC Progressively Aligns Chromosomal Arms in Caulobacter crescentus but Is Antagonized by Convergent Transcription. Cell Rep. 20, 2057–2071 (2017).

40. G. A. Busslinger, R. R. Stocsits, P. van der Lelij, E. Axelsson, A. Tedeschi, N. Galjart, J.-M. Peters, Cohesin is positioned in mammalian genomes by transcription, CTCF and Wapl. Nature. 544, 503–507 (2017).

41. K. Maeshima, S. Ide, M. Babokhov, Dynamic chromatin organization without the 30-nm fiber. Curr. Opin. Cell Biol. 58, 95–104 (2019).

42. V. I. Risca, S. K. Denny, A. F. Straight, W. J. Greenleaf, Variable chromatin structure revealed by in situ spatially correlated DNA cleavage mapping. Nature. 541, 237–241 (2017).

43. K. R. Kieffer-Kwon, K. Nimura, S. S. P. Rao, J. Xu, S. Jung, A. Pekowska, M. Dose, E. Stevens, E. Mathe, P. Dong, S. C. Huang, M. A. Ricci, L. Baranello, Y. Zheng, F. T. Ardori, W. Resch, D. Stavreva, S. Nelson, M. McAndrew, A. Casellas, E. Finn, C. Gregory, B. G. St. Hilaire, S. M. Johnson, W. Dubois, M. P. Cosma, E. Batchelor, D. Levens, R. D. Phair, T. Misteli, L. Tessarollo, G. Hager, M. Lakadamyali, Z. Liu, M. Floer, H. Shroff, E. L. Aiden, R. Casellas, Myc Regulates Chromatin Decompaction and Nuclear Architecture during B Cell Activation. Mol. Cell. 67, 566–578.e10 (2017).

44. T. H. S. Hsieh, C. Cattoglio, E. Slobodyanyuk, A. S. Hansen, O. J. Rando, R. Tjian, X. Darzacq, Resolving the 3D Landscape of Transcription-Linked Mammalian Chromatin Folding. Mol. Cell. 78, 539–553.e8 (2020).

45. M. M. Spain, S. G. Swygert, T. Tsukiyama, Preparation and analysis of saccharomyces cerevisiae quiescent cells in Cellular Quiescence. Methods in Molecular Biology (Humana Press, New York, NY, vol. 1686, 2018), pp. 125–135.

46. J. Schindelin, I. Arganda-Carreras, E. Frise, V. Kaynig, M. Longair, T. Pietzsch, S. Preibisch, C. Rueden, S. Saalfeld, B. Schmid, J.-Y. Tinevez, D. J. White, V. Hartenstein, K. Eliceiri, P. Tomancak, A. Cardona, Fiji: an open-source platform for biological-image analysis. Nat. Methods. 9, 676–682 (2012).

47. T. H. Giddings, E. T. O’Toole, M. Morphew, D. N. Mastronarde, J. R. McIntosh, M. Winey, Using rapid freeze and freeze-substitution for the preparation of yeast cells for electron microscopy and three-dimensional analysis. Methods Cell Biol. 67, 27–42 (2001).

48. D. N. Mastronarde, Dual-axis tomography: an approach with alignment methods that preserve resolution. J. Struct. Biol. 120, 343–352 (1997).

49. G. Arya, T. Schlick, A tale of tails: how histone tails mediate chromatin compaction in different salt and linker histone environments. J. Phys. Chem. A. 113, 4045–4059 (2009).

50. O. Perišić, R. Collepardo-Guevara, T. Schlick, Modeling studies of chromatin fiber structure as a function of DNA linker length. J. Mol. Biol. 403, 777–802 (2010).

51. D. A. Beard, T. Schlick, Modeling salt-mediated electrostatics of macromolecules: the discrete surface charge optimization algorithm and its application to the nucleosome. Biopolymers. 58, 106–15 (2001).

52. H. Jian, A. V. Vologodskii, T. Schlick, A combined wormlike-chain and bead model for dynamic simulations of long linear DNA. J. Comput. Phys. 136, 168–179 (1997).

53. G. Arya, Q. Zhang, T. Schlick, Flexible histone tails in a new mesoscopic oligonucleosome model. Biophys. J. 91, 133–150 (2006).

54. A. Luque, R. Collepardo-Guevara, S. Grigoryev, T. Schlick, Dynamic condensation of linker histone C-terminal domain regulates chromatin structure. Nucleic Acids Res. 42, 7553–7560 (2014).

55. O. Perisic, S. Portillo-Ledesma, T. Schlick, Sensitive effect of linker histone binding mode and subtype on chromatin condensation. Nucleic Acids Res. 47, 4948–4957 (2019).

56. R. Collepardo-Guevara, G. Portella, M. Vendruscolo, D. Frenkel, T. Schlick, M. Orozco, Chromatin unfolding by epigenetic modifications explained by dramatic impairment of internucleosome interactions: A multiscale computational study. J. Am. Chem. Soc. 137, 10205–10215 (2015).

57. C. G. Baumann, S. B. Smith, V. A. Bloomfield, C. Bustamante, Ionic effects on the elasticity of single DNA molecules. Proc. Natl. Acad. Sci. U. S. A. 94, 6185–6190 (1997).

58. K. Chen, Y. Xi, X. Pan, Z. Li, K. Kaestner, J. Tyler, S. Dent, X. He, W. Li, DANPOS: Dynamic analysis of nucleosome position and occupancy by sequencing. Genome Res. 23, 341–351 (2013).

59. Y. Zhang, T. Liu, C. A. Meyer, J. Eeckhoute, D. S. Johnson, B. E. Bernstein, C. Nussbaum, R. M. Myers, M. Brown, W. Li, X. S. Shirley, Model-based analysis of ChIP-Seq (MACS). Genome Biol. 9, R137 (2008).

60. H. R. Drew, A. A. Travers, DNA bending and its relation to nucleosome positioning. J. Mol. Biol. 186, 773–790 (1985).

61. N. Metropolis, S. Ulam, The Monte Carlo Method. J. Am. Stat. Assoc. 44, 335 (1949).

62. M. N. Rosenbluth, A. W. Rosenbluth, Monte carlo calculation of the average extension of molecular chains. J. Chem. Phys. 23, 356–359 (1955).

63. P. J. G. Butler, J. O. Thomas, Dinucleosomes show compaction by ionic strength, consistent with bending of linker DNA. J. Mol. Biol. 281, 401–407 (1998).

64. B. Langmead, S. L. Salzberg, Fast gapped-read alignment with Bowtie 2. Nat. Methods. 9, 357–359 (2012).

65. N. C. Durand, J. T. Robinson, M. S. Shamim, I. Machol, J. P. Mesirov, E. S. Lander, E. L. Aiden, Juicebox Provides a Visualization System for Hi-C Contact Maps with Unlimited Zoom. Cell Syst. 3, 99–101 (2016).

66. L. Giorgetti, B. R. Lajoie, A. C. Carter, M. Attia, Y. Zhan, J. Xu, C. J. Chen, N. Kaplan, H. Y. Chang, E. Heard, J. Dekker, Structural organization of the inactive X chromosome in the mouse. Nature. 535, 575–579 (2016).

67. H. Li, B. Handsaker, A. Wysoker, T. Fennell, J. Ruan, N. Homer, G. Marth, G. Abecasis, R. Durbin, The Sequence Alignment/Map format and SAMtools. Bioinformatics. 25, 2078–2079 (2009).

68. F. Ramírez, D. P. Ryan, B. Grüning, V. Bhardwaj, F. Kilpert, A. S. Richter, S. Heyne, F. Dündar, T. Manke, deepTools2: A next generation web server for deep-sequencing data analysis. Nucleic Acids Res. 44, W160–W165 (2016).

69. N. H. Freese, D. C. Norris, A. E. Loraine, Integrated genome browser: Visual analytics platform for genomics. Bioinformatics. 32, 2089–2095 (2016).

